# Single-molecule nanopore sequencing reveals spatial coordination of rRNA modifications in human ribosomes

**DOI:** 10.64898/2026.09.11.751006

**Authors:** George Ettenger, Aaron M. Fleming

**Affiliations:** Department of Chemistry, University of Utah, 315 S 1400 E, Salt Lake City, UT 84112-0850

## Abstract

Ribosomal RNA has a high density of epitranscriptomic modifications essential for faithful translation, and their levels vary across cell types and in disease. Typically, rRNA modifications are quantified in bulk, and therefore, coordination on individual RNA molecules has remained poorly understood. Herein, modification-aware, single-molecule nanopore sequencing enabled detection of the co-occurrence of rRNA modifications on individual transcripts, resolving coordination invisible to ensemble methods. An analytical framework was established that separates co-occurrence from read-quality, false-positive, calling-artifact, near-saturation, and global modification-level confounds, using human rRNAs from HEK293T cells as the test dataset. The method was applied to four human cell lines to find that co-occurrence is not generally explained by a shared small nucleolar RNA (snoRNA) guide; instead, coordinated modifications cluster locally, within ∼100 nucleotides and 35-40 Å in the folded ribosome. For one shared-guide pair, coordination increased as guide levels fell across cell lines, possibly indicating an all-or-nothing mode of modification per molecule under limiting guide availability. Further, the results revealed rRNA heterogeneity between the cells in overall modification levels. Finally, levofloxacin remodeled specific modification sites and their local co-occurrence networks, showing that the approach can reveal small-molecule perturbation of rRNA modification networks.

## Introduction

Ribosomal RNA is a central component of the ribosome, a ∼4 MDa biological machine that translates the genetic code in mRNA into protein. A feature that enables efficient, high-fidelity translation is a set of rRNA chemical modifications decorating the bases and sugars of specific nucleotides in the 5.8S, 18S, and 28S rRNAs of human cytosolic ribosomes [1]. Structural studies place many of these modifications at functionally important regions of the ribosome [2,3], where they tune the ribosome structure required for accurate codon decoding and peptide-bond formation [4]. Prior studies identified that rRNA modifications differ between cell types and in diseased tissues [1,5]; however, it remains open whether the observed changes reflect independent cellular events or coordinated modification programs.

The ribosome is often taught as a static, protein-producing machine. However, many observations support an alternative view in which ribosomes are heterogeneous and may themselves contribute to cellular regulation. Ribosomal proteins are not uniformly expressed across cell types, producing compositional differences between cells [6,7]. The human genome carries hundreds to thousands of rDNA copies that are not identical in sequence, introducing rRNA sequence variants into the pool of ribosomes [8–10]. Furthermore, the 5.8S, 18S, and 28S rRNAs of human cytosolic ribosomes carry ∼250 chemical modifications spanning 15 distinct chemistries (Figures 1A and S1) [1–3,5,11,12]. Quantitative mass spectrometry mapping has shown that while many of these modifications are installed at near-stoichiometric levels (>90% modified), 34% are sub-stoichiometric in their abundance (<90% modified) [8], and it is this partial modification that creates the ribosomal rRNA modification heterogeneity. Together, these sources of variation motivate the specialized-ribosome hypothesis, in which the ribosome sub-populations preferentially translate distinct classes of mRNA [13–15]. Testing whether rRNA modifications form networks at the level of molecules as part of a dynamic chemical layer of ribosome specialization is information that bulk measurements like mass spectrometry cannot provide.

**Figure 1.**
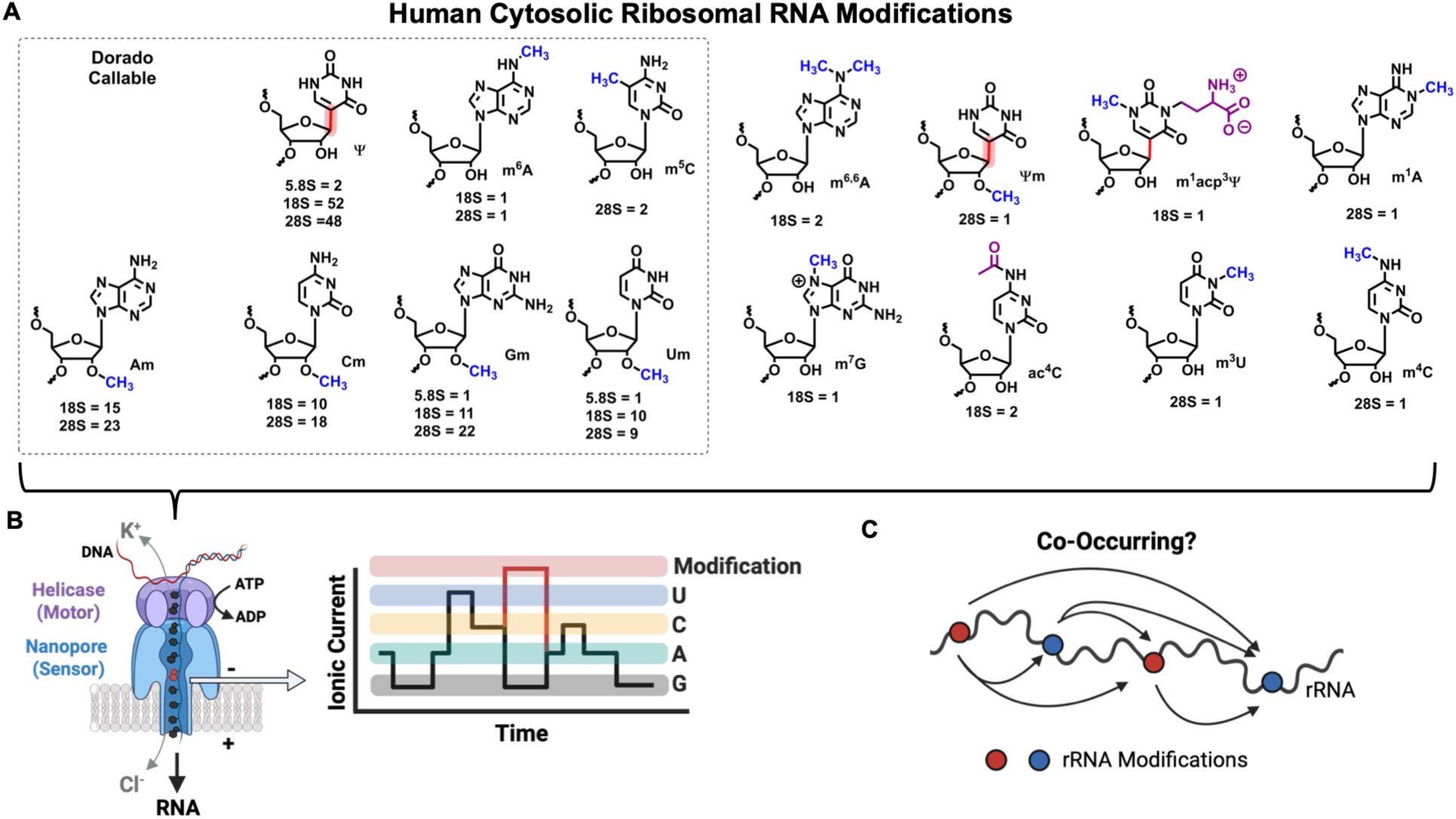
Ribosomal RNA is decorated with chemical modifications that can be analyzed by nanopore sequencing to determine modification pair co-occurrence. (A) The 15 different rRNA modification structures and their counts in the 5.8S, 18S, and 28S human rRNAs. The modifications called by Dorado v5.2 (https://github.com/nanoporetech/dorado/) are shown in the dashed box. (B) Direct RNA nanopore sequencing enables modification analysis. (C) By analyzing populations of sequenced single molecules of rRNA, the co-occurrence of the modifications can be measured.

Current approaches for identifying rRNA modification sites include structural methods, mass spectrometry, and quantitative sequencing [1–3,11,12,16–18]. Structural methods (cryo-EM and X-ray crystallography) resolve the three-dimensional positions of modifications within a ribosome, revealing their clustering at functional centers; however, they report an ensemble average over the population of particles analyzed and require resolution sufficient to discern modifications. Mass spectrometry is generally used to quantify modification occupancy by analyzing nucleosides digested from RNA polymers, resulting in loss of sequence-context information [19]. Developments in RNA modification analysis on intact strands can address where modifications change without complete digestion [20,21], but the results are still obtained as ensemble averages. Quantitative sequencing approaches, such as RiboMeth-seq for 2′-*O*-methylation (Nm) and bisulfite sequencing for pseudouridine (Ψ), provide site-resolved occupancy levels in rRNA strands [16–18,22]; however, their accuracy depends on the efficiency of the modification-specific chemical step on native RNA [23]. Mass spectrometry and quantitative sequencing report how much of a modification is present across a population of rRNAs, but neither can determine whether modifications co-occur on the same molecule.

Nanopore direct RNA sequencing can detect multiple chemical modifications along individual RNA molecules sequenced end to end [5,24–27], providing a tool to answer questions about rRNA modification heterogeneity at the molecule level that ensemble methods cannot address (Figure 1b). Advances have improved the sequencer and the computational tools for modification-aware RNA sequencing [28–30]. The sequencer enables analysis of single ionic-current versus time traces with a neural-network base caller, providing reads on individual molecules without restriction on transcript length and yielding both sequence and modification state (Figure 1b). Challenges remain in sequencing with quantitative accuracy and complete modification awareness. Nonetheless, the feasibility of extracting modification co-occurrence from such reads has been demonstrated on cellular RNA using custom-developed data-analysis tools [31–33]. What remains lacking is an approach suited to rRNA, whose high modification density, mixture of base and sugar modifications, and characteristic calling artifacts have been controlled for to reveal modification co-occurrence on single molecules; thus, we built our approach on a standard commercial base caller (Dorado, v5.2; Figure 1a), allowing portability to any laboratory performing nanopore sequencing and enabling incorporation of future Dorado-supported modification models.

The present work presents a method for measuring rRNA modification co-occurrence by analyzing many rRNA molecules one at a time, using Oxford Nanopore Technologies (ONT) modification-aware Dorado base callers to detect m^6^A, m^5^C, Ψ, and the four 2′-*O*-methylated nucleotides; these modifications account for >95% of modified positions in human rRNA (Figure 1a) [1]. The method identifies and controls for artifacts that confound single-molecule modification analysis, such as read quality, modification clustering, and global strand-to-strand modification levels. The approach was deployed on the 5.8S, 18S, and 28S rRNAs across four human cell lines, revealing modification co-occurrence networks in the 18S and 28S rRNA that can differ between cells. We find that co-occurrence is generally organized by proximity, clustering among modifications close in sequence and in three-dimensional space rather than among those sharing a common snoRNA guide; however, for one shared-guide pair the co-occurrence tracked inversely with snoRNA levels across the cells studied, suggesting modification writing may be all-or-nothing when a guide is limiting. Lastly, a small molecule was used to selectively edit methyl groups on the 18S rRNA, leading to changes in the rRNA modification co-occurrence networks involving the edited sites.

## Material and Methods

### Cell culture and RNA extraction

HEK293T, HeLa, MCF7, and HCT116 cells were grown in DMEM supplemented with 10% FBS, 1x GlutaMAX™, 1x non-essential amino acids, and 1x pen/strep supplement. The cells were maintained in a cell culture incubator under 5% CO₂ with 80% humidity at 37 °C. HEK293T cells were exposed to 25 µM levofloxacin (Lev) for 48 h following a literature protocol. Once cells reached 80% confluency, they were harvested by centrifugation. All cell pellets were stored at −80 °C until studied. Total RNA was extracted using the Zymo RNA extraction kit following the manufacturer’s protocol. RNA integrity was assessed by agarose gel electrophoresis. Before library preparation, 2 μg of total RNA was 3′ poly-A tailed using a commercial poly-A tailing kit (NEB) following the manufacturer’s instructions.

### Quantification of snoRNAs by RT-qPCR

The SNORD50A and SNORD50B snoRNA expression profiles were measured using the Luna Universal One-Step RT-qPCR Kit (NEB), following the manufacturer’s protocol. The primer sequences were previously reported [34]: SNORD50A forward, 5′-TAT CTG TGA TGA TCT TAT CCC GAA CCT GAA C-3′ and reverse, 5′-ATC TCA GAA GCC AGA TCC GTA A-3′; SNORD50B forward, 5′-TAA TCA ATG ATG AAA CCT ATC CCG-3′ and reverse, 5′-GCT TCG GCA GCA CAT ATA CTA AAA T-3′. The U6 snRNA was used as an internal control with the primer pair 5′-GCT TCG GCA GCA CAT ATA TAC TAA AAA AT -3′ and 5′-CGC TTC ACG AAT TTG CGT GTC AT -3′. Total RNA for the RT-qPCR analysis was obtained as described above. The RT-qPCR C_T_ values were analyzed using the 2^(−ΔΔC_T_) method [35], with expression normalized to the U6 snRNA for the plots presented.

### Nanopore sequencing and basecalling

The purified, poly-A-tailed RNA was used for library preparation with the SQK-RNA004 direct RNA sequencing kit (ONT) following the manufacturer’s protocol. The library was loaded onto an RNA flow cell (ONT) and sequenced per the manufacturer’s protocol, with passed reads having a Q > 8. The ionic current versus time traces were saved in POD5 format. Passed reads were basecalled with Dorado (v5.2) using the modification-aware models at super-accuracy (sup) setting (rna004_130bps_sup@v5.2.0) running the default parameters. Because Dorado permits only one modification model per canonical base per pass, each POD5 dataset was basecalled four times using the four available models: (1) m6A and Am; (2) m5C and Cm; (3) Gm; and (4) Ψ and Um. All four BAM files contain the same reads, identified by a common read_id, permitting per-read joins across modification classes. Reads were aligned to an rRNA reference containing the 5.8S, 18S, and 28S human cytosolic rRNA sequences using minimap2 within Dorado, with the preset RNA alignment parameters. Alignments were sorted and indexed with samtools (v1.21).

### Per-read modification calls

Per-read modification states were extracted with modkit (https://github.com/nanoporetech/modkit) using extract calls with the filter-threshold set to 0.7, restricted to the known modified sites on each rRNA (--include-bed) and to the relevant reference sequence (--region). For each read–site pair, the retained fields were read_id, ref_position, call_prob, call_code, and fail. Critically, each site was scored by its own modification code, not by any modification call at that position. A site annotated as Um, for example, was scored as modified only if its call code corresponded to Um, preventing cross-modification bleed-through into the data.

Because the four modification-annotated BAM files were derived from the same POD5 files, the per-read modification states were joined across the four files by read_id to construct a binary read (r) x site (s) matrix M, where M(r,s) = 1 if read r carried the annotated modification at site s, 0 if it was called unmodified, and missing if read r did not span site s or its call failed the threshold. Analyzable sites were defined as the intersection of three criteria: (1) sites were restricted to previously reported modifications in the human 5.8S, 18S, and 28S rRNA [1–3,11,12]; (2) only modifications with a corresponding Dorado model were retained (m^6^A, m^5^C, Ψ, Am, Cm, Gm, Um), while modifications without a model (e.g., ac^4^C, m^7^G, m^1^A, m^3^U, m^1^acp^3^Ψ,and Ψm) were not analyzed, and (3) sites were required to exceed a minimum modification fraction and read depth (≥25% and ≥100 reads) to be analyzed, since sites called at low fractional levels near the background noise level despite high coverage do not carry per-read variance and cannot contribute to a co-occurrence estimate. The site table annotating the rRNA modifications analyzed and their positions is provided in Figure S1.

### Co-occurrence statistics

For each pair of modified sites (i, j) within an rRNA, co-occurrence was quantified on the reads spanning both sites. The 2 x 2 contingency table of joint modification states (both modified / i only / j only / neither) yields an odds ratio (OR):

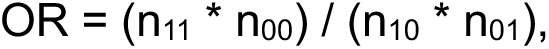

where OR > 1 indicates that modifications at the two sites co-occur on the same molecule more often than expected under independence, OR = 1 indicates independence, and OR < 1 indicates mutual exclusivity.

### Adjustment for read quality and global modification level

Read quality could in principle drive spurious correlation, because a high-quality read may yield confident modification calls at both sites, generating co-occurrence with no biological basis. To control for this, OR values were computed with the Mantel–Haenszel (MH) estimator [36] stratifying reads into five strata by alignment identity (aln_identity) and, by mean read quality (mean_q). The MH estimator pools the stratum-specific odds ratios:

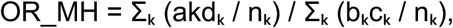

where aₖ = reads modified at both sites, bₖ = modified at i only, cₖ = modified at j only, dₖ = neither modified, and nₖ = the stratum total.

To remove the possible confound that global modification level per strand could impact the results, a second stratification on the OR calculation was applied based on global modification level (G). For each read, the fraction of covered variance-rich sites that were modified, where variance-rich sites are those with a marginal modification frequency between 0.25 and 0.90, was calculated. Reads were binned into five strata of G, and the OR values for every site pair was recomputed pooled across the strata to yield an adjusted value (i.e., OR_adj). Unless otherwise noted, reported co-occurrences are those that remain significant after this global-level adjustment (OR_adj); pairs called on the raw association but lost after adjustment were flagged as possibly globally confounded.

Statistical analysis of the MH odds ratio, was calculated with the Cochran–Mantel– Haenszel (CMH) χ² statistic [36], and the Robins–Breslow–Greenland (RBG) 95% confidence intervals were computed with a custom Python implementation [37]; χ² statistics were converted to P-values using SciPy (v1.13.1). P-values were corrected for multiple testing across all site pairs within an rRNA using the Benjamini–Hochberg (BH) false-discovery-rate (FDR) procedure [38]. A pair was therefore called only if it satisfied both a significance and an effect-size criterion: FDR q < 0.01 and OR_adj ≥ 1.3 (positive co-occurrence) or FDR q < 0.01 and OR_adj ≤ 0.77 (mutual exclusivity).

### Software and availability

All analyses were implemented in Python 3.12.7 using numpy 1.26.4, pandas 2.2.2, scipy 1.13.1, and matplotlib 3.9.2. Commercial LLMs assisted in generation of the Python code.

## Results

### Developing a tool for rRNA modification co-occurrence analysis

The approach to measuring co-occurrence between rRNA epitranscriptomic modifications on individual nanopore reads was applied to modification-aware basecalls from Dorado (v5.2). Basecalling of nanopore data with Dorado, exhibits a broad range of accuracies across the modifications it has been trained on that occur in rRNA (non-DRACH m^6^A, m^5^C, Ψ, Am, Cm, Gm, and Um; Figure 1a) [39]. Several situations are known to challenge modification calling, including local sequence contexts that yield poor ionic-current separation between modified and canonical strands [26,40,41]; clustering of modifications [42], which confounds attributing the ionic-current change to a single position; modifications present at low occupancy; and outright false-positive calls [39,41,43]. Each of these was controlled for, filtered, or minimized by the method. As an advantage for the analysis, these errors are largely systematic, resulting in good reproducibility between replicate analyses, as described below.

Method development was achieved on direct RNA nanopore sequencing data for rRNA from HEK293T cells [44]. From the same POD5 file, basecalling was performed at the super-accuracy (sup) level with modification-aware Dorado models applied in four passes. Each pass was applied to one of the four canonical nucleotides at a time, including m^6^A with Am, m^5^C with Cm, Gm alone, and Ψ with Um (Figure 2a). Because the four passes derive from identical raw signal, every read appears in all four datasets under a common read_id, permitting a per-molecule join across the modification classes. Analysis was restricted to a curated list of human rRNA modification sites established by cryo-EM and mass spectrometry (Figure S1) [1–3,11,12], an intersection that excludes false-positive calls at unannotated sites. The necessity of this step is illustrated by the m^5^C model as the most extreme example, which called 44 false positives in the 28S rRNA (>0.5 fractional occupancy and >100x coverage) that are absent from the annotated list (Figure 1a). From the only two curated 28S m^5^C sites, one was found above the threshold values for modification analysis. Only rRNA modifications for which a model exists were included in the analysis; modifications such as 18S m^1^acp^3^Ψ1248 were not inspected (Figure 1a). Each site was scored by its own modification code, preventing cross-modification bleed-through (e.g., an m^6^A call at an annotated Am site) from being counted as evidence of modification.

**Figure 2.**
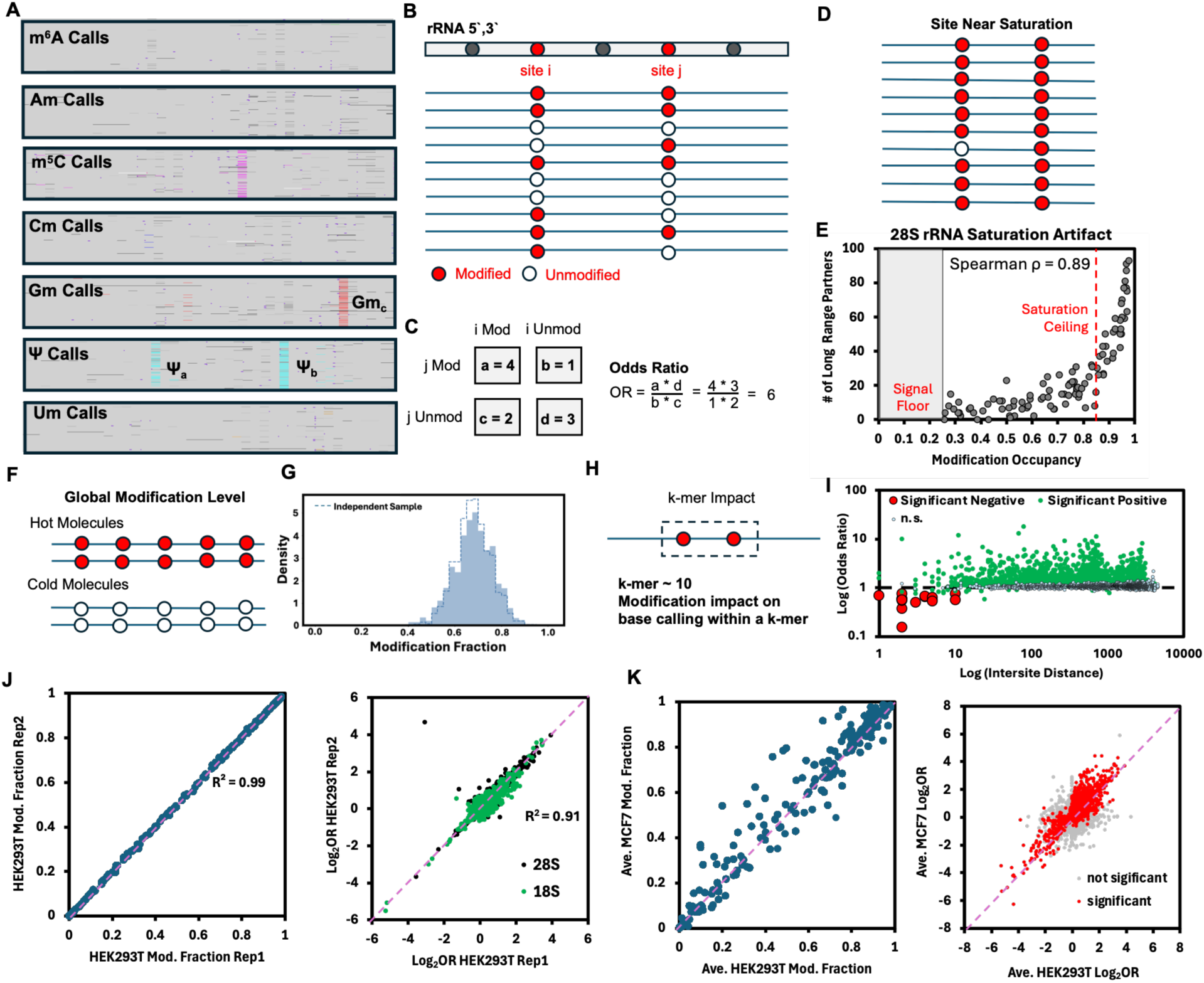
Analysis to control for confounding factors in the co-occurrence analysis of rRNA modifications on individual molecules. (A) Single-molecule reads for each of the modification-aware models for a section of the 5.8S rRNA from HEK293T cells; each panel represents ∼100 transcripts. (B) Example single-read data for co-occurrence analysis and (C) the corresponding odds-ratio calculation. (D) Example single-read data illustrating that modification saturation (E) generates hubs of co-occurrence that are difficult to distinguish from a mathematical artifact. (F) The global modification level of rRNA is (G) heterogeneous and was controlled for in the odds-ratio analysis. (H) When two rRNA modifications reside within a common ∼10-nucleotide k-mer, (I) significant mutual exclusion was observed, leading us to exclude co-occurrence analysis within the k-mer window. Data in the plots for pipeline development were obtained from 5,152 28S rRNA modification pairs with depths ranging from 1,009 to 65,404. Reproducibility plots of modification fraction measurements and OR analysis for (J) HEK293T biological replicates and (K) differences in average values between HEK293T and MCF7 cells. Data for the replicate analyses were obtained from the 18S and 28S rRNA, with 14,402 modification pairs with depth ranging from 1,681 to 67,164.

Using read_id, the four datasets were joined per molecule to construct a binary read x site matrix, in which each entry records whether a read carried the annotated modification at a site (1), was called canonical (0), or did not span the site (blank; Figure 2b). For each site pair, the reads spanning both sites populate a 2 x 2 contingency table from which an odds ratio (OR) was computed (Figure 2c). An OR > 1 indicates that modifications at the two sites co-occur on the same molecule more often than expected under independence, OR = 1 indicates independence, and OR < 1 indicates mutual exclusivity (Figure 2b; Figures S2 and S3 for summary values of the analysis).

Significance was assessed by the Cochran–Mantel–Haenszel χ^2^ test, with the Benjamini– Hochberg FDR procedure correcting for multiple testing across all site pairs sequenced within an rRNA (theoretical maximum values: 28S rRNA, n = 9,180 pairs; 18S rRNA, n = 5,253 pairs; 5.8S rRNA, n = 6 pairs). Because per-pair sequencing depth is very high (>10^3^), essentially any departure from OR = 1 attains statistical significance, so P-values alone are uninformative at this depth. Pairs were consequently called only when they satisfied both a significance and an effect-size criterion with an FDR q < 0.01 and OR ≥ 1.3 (positive co-occurrence), or FDR q < 0.01 and OR ≤ 0.77 (mutual exclusivity).

We controlled for several artifacts in the data. First, nanopore data are known to be error-prone, which impacts the per-read quality score. Reads were stratified by read quality using the MH estimator to recompute the OR values. Stratification on mean read quality yielded estimates concordant with those calculated without read-quality stratification (median difference = 0.56%; Figure S4). Second, modification level, whether high or low, can distort the OR calculation, because rare joint states at extreme occupancies produce mathematically unstable OR values (Figures 2d and S5). This was evident when the number of co-occurring partners for each 28S rRNA modification was plotted against its modification level, which revealed a positive correlation (Figure 2e, ρ = 0.89). Therefore, the analysis window was bounded with a floor of 0.25 and a ceiling of 0.85; the floor keeps sequencing background noise from dominating measurements at lowly modified sites, while the ceiling excludes near-saturated sites, which carry too few unmodified reads to yield a stable odds ratio and consequently emerge as spurious hubs. As a result of modification levels differing between cell lines, these thresholds retained different numbers of sites in each cell; for example, in HEK293T cells, they retained 81 of 134 analyzable 28S rRNA sites (61%) and 73 of 103 analyzable 18S rRNA sites (71%; Figure S3).

The most impactful confound handled molecules that differ in how heavily they are modified overall, such that two sites can appear to co-occur simply by tracking the global modification level rather than through any specific relationship (Figure 2f). Stratifying reads by their per-molecule global modification level with the MH estimator removed 53% of the raw co-occurrences, making this the single largest correction applied (Figure S5). Left uncorrected, this confound is sufficient to make the modifications appear as a coordinated network, in which many pairs acquire a positive association and tightly interconnected hubs and communities emerge when lightly and heavily modified molecules are compared.

To assess whether these global modification states are themselves coordinated, the observed per-molecule modification-level distribution was compared to a randomized distribution generated by shuffling each site’s modification state across molecules while preserving its marginal frequency, which breaks any per-molecule coupling (Figure 2g). The observed distribution was 1.2x broader than the randomized one (Figure 2g, dotted distribution), indicating that the modification states of the rRNA are not fully independent (Figure S6). Thus, single-transcript sequencing revealed heterogeneous global modification states among ribosomes, and this global modification propensity accounts for a substantial fraction of the observed pairwise co-occurrence. The co-occurrences lost upon global-level stratification were predominantly those involving sites near the ceiling threshold, whereas pairs at intermediate modification fraction were retained (Figure S6). These retained co-occurrences cannot be explained by global modification level, read-quality differences, or near-saturation, supporting a biological basis for their co-occurrence. Moreover, the persistence of a broadened global modification-level distribution after controlling for nanopore artifacts points to possible per-molecule differences in rRNA biogenesis; this is considered in the Discussion but is not a focus of the present work.

As a consequence of the nanopore sequencer measuring ionic current from a k-mer of nucleotides (∼10 nt) rather than a single base, a modification at one site can suppress the call of another neighbor modification, producing a false appearance of mutual exclusivity (Figures 2h/i and S5); negative associations were interpreted as biologically relevant only for sites more than 10 nt apart. In the remainder of the text, reported OR values are those adjusted with the MH estimator for both read quality and global modification level (OR_adj). In the text and figures, OR_adj is simply referred to as OR, as this is the only parameter used to discuss the data. The OR values were log_2_-transformed to maintain symmetry in the increase or decrease from 1. For example, OR = 0.5 or 2 are a 2-fold reduction or increase from 1, respectively, while their magnitudes from 1 differ; in contrast, log_2_ transformation of 0.5 or 2 equals -1 and 1, respectively, having the same magnitude of change from 1, consistent with their fold change; thus, an OR ≥ 1.3 is log_2_OR ≥ 0.38 and an OR ≤ 0.78 is a log_2_OR ≤ -0.38. Lastly, the reproducibility in modification fraction and log_2_OR analyses between HEK293T biological replicates demonstrates reproducibility in the nanopore data, even though it is less accurate (Figure 2j). The average modification fraction and log_2_OR values between HEK293T and MCF7 analyses illustrate the differences observed between the cell lines that form the basis of cell line differences in rRNA modification co-occurrence (Figure 2k).

#### rRNA modification co-occurrence has spatial dependency

Having established a confound-controlled measure of co-occurrence, we asked how coordinated modifications are distributed across the 18S and 28S rRNAs; the data for the 5.8S rRNA from HEK293T cells are discussed in Figure S7, which turned out to be an illustrative case for adjusting the data to remove confounds. Data from all four cell lines were aggregated and analyzed globally for trends in the dependence of positive co-occurrence on sequence (2D) and three-dimensional (3D) distance (2-20% significant pairs per strand from the four cell lines; Figure S3). Modification pairs reaching significant positive co-occurrence (q < 0.01 and log_2_OR ≥ 0.38) were binned by their separation in sequence space or in 3D structure space in the human ribosome and plotted as the fraction of significant pairs per distance bin (Figure 3a, red lines). To avoid modification level as a confounder, only pairs in which both sites had modification fraction >0.25 and at least one site with modification fraction <0.85, were included (Figure 2e). Null distributions were generated by randomizing the data from all strands 1,000 times and reanalyzing the distance dependence (Figure 3a, gray area).

**Figure 3.**
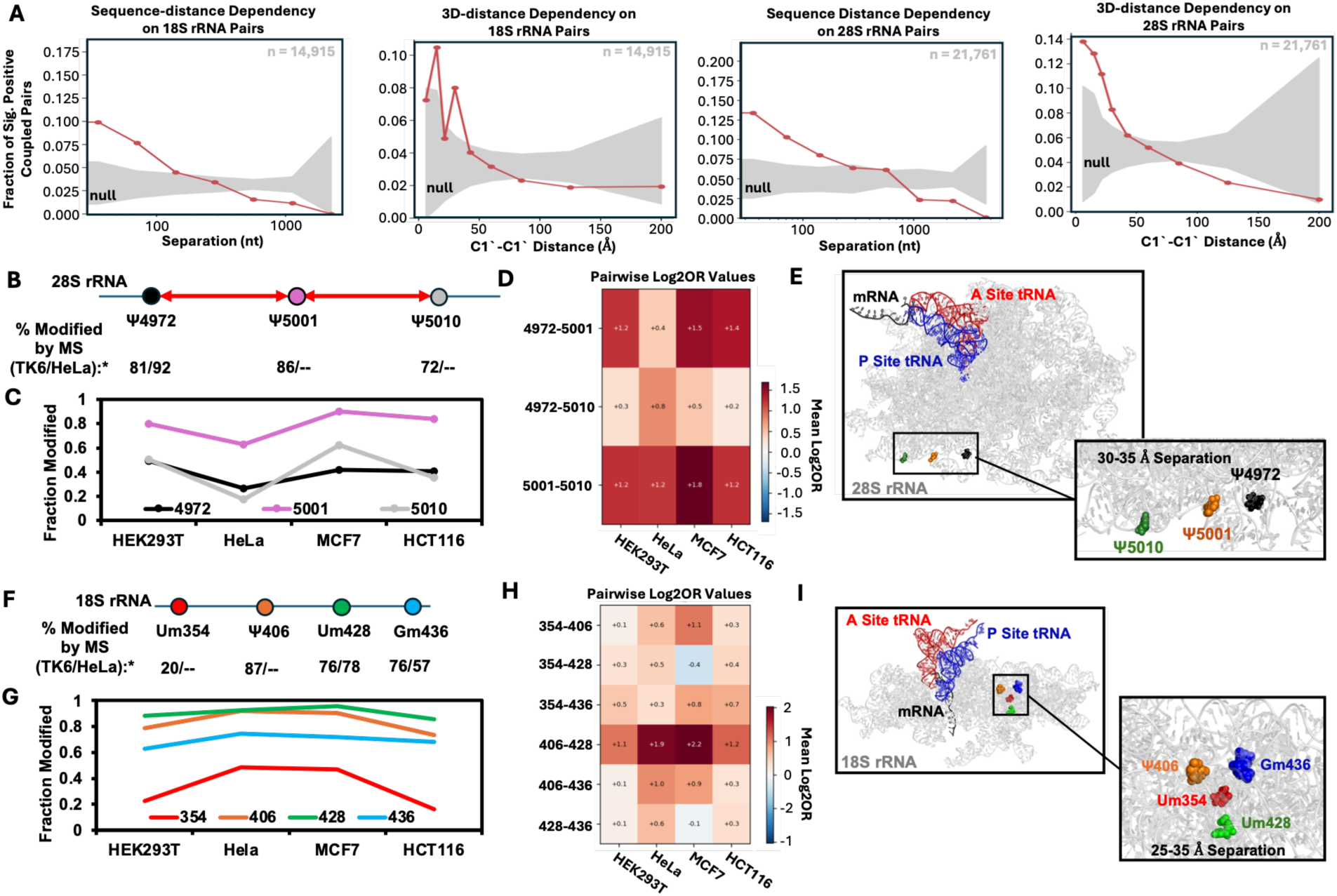
Co-occurrence analysis of rRNA modifications on individual transcripts identifies spatial trends and local networks. (A) Dependence of the fraction of statistically significant co-occurring pairs on sequence (2D) and three-dimensional (3D) distance in the 18S and 28S rRNAs, aggregated across HEK293T, HeLa, MCF7, and HCT116 cells. Pairs were included when both sites had an occupancy > 0.25 and at least one site had an occupancy < 0.85. (B) A Ψ network in the 28S rRNA at positions 4972, 5001, and 5010; *these sites were previously found by mass spectrometry to be written sub-stoichiometrically in TK6 and HeLa rRNA [1]. (C) Nanopore-measured modification levels across the four cell lines and (D) the co-occurrence log_2_OR values for the three-site network. (E) Structural map of the three Ψ residues on the human 28S rRNA (PDB 6Y0G), shown with the mRNA and A-and P-site tRNAs to illustrate that the network is distal to the peptidyl transferase center. (F) A four-site co-occurring network in the 18S rRNA (Um354, Ψ406, Um428, Gm436), *whose members were previously found by mass spectrometry to be written sub-stoichiometrically in TK6 and HeLa cells [1]. (G) Nanopore-measured modification levels across the four cell lines and (H) the log_2_OR values showing favorable co-occurrence. (I) Structural maps for the modifications are shown in PDB 6Y0G [45], which includes the mRNA and A- and P-site tRNAs to illustrate the decoding center; all reported C1′–C1′ distances were measured on PDB 8QO1 [3].

In both the 18S and 28S rRNA, a greater fraction of significant, positive co-occurring pairs than in cell lines compared to the null (independent) dataset was found at sequence distances less than ∼100 nt (Figure 3a). The 3D distances were obtained from the C1′–C1′ measurements between every pair of rRNA modification sites in a reported human ribosome structure (pdb: 8QO1 [3]). In the 18S rRNA, the trend was not monotonic at the shortest distances but was generally greater than the null sample out to ∼35 Å of separation (Figure 3a). In the 28S rRNA, co-occurrence in the cell samples exceeded the null sample out to nearly 40 Å (Figure 3a). Together, these findings indicate that co-occurrence is favored for modifications within ∼100 nucleotides of sequence and within ∼35–40 Å of 3D space, depending on the rRNA sequence.

Within the aggregated data, two networks were examined in detail. The first, in the 28S rRNA, comprised the three Ψ residues at positions 4972, 5001, and 5010, which formed a positive, significant network (q < 0.01 and OR ≥ 1.3, i.e., log_2_OR ≥ 0.38; Figure 3b). Each site reached a nanopore modification fraction within the 0.25–0.85 window in at least one of the four cell lines (Figure 3c), and prior mass spectrometry results indicate these sites are sub-stoichiometric in TK6 or HeLa cells [1], permitting co-occurrence to be measured on individual molecules (Figure 3c, asterisks). The log_2_OR values support a network among all three residues, with co-occurrence decreasing in the order Ψ5001–Ψ5010 > Ψ4972–Ψ5001 > Ψ4972–Ψ5010 (Figure 3d). MCF7 cells showed the largest log_2_OR values; however, the three sites approach saturation in MCF7, so these elevated values likely reflect near-saturation inflation of the OR value rather than stronger biological coupling; they are interpreted with caution. In three dimensions, the three C1‘-C1’ atoms in the co-occurring network residues reside within 35 Å of one another, and the network is distal to the peptidyl transferase center, as illustrated with the location of the mRNA, A-and P-site tRNAs in the structure (Figure 3e).

The second network was comprised of 18S Um354, Ψ406, Um428, and Gm436. Prior mass spectrometry found all four modifications to be sub-stoichiometric in TK6 and HeLa cells [39], with Um354 present at only ∼20% occupancy (Figure 3f); thus, a privileged subset of rRNA molecules carrying these modifications is plausible. Across the four cell lines, Um354 was most highly modified in HeLa and MCF7 cells and least modified in HEK293T and HCT116 cells (Figure 3g). Um354 co-occurrence with each of the other three sites exceeded the significance threshold (q < 0.01 and log_2_OR ≥ 0.38) in the four cell lines, with one exception being Um354–Um428 showing mutual exclusivity in MCF7 cells (Figure 3h). The Ψ406–Um428 edge anchored the network with the largest log_2_OR; however, these two sites are the most highly modified, and this edge may be partly inflated by near-saturation and is interpreted with caution. Critically, Um354, which is the least-modified site, remained coupled to the other three modifications, albeit at lower log_2_OR values (Figure 3h). For Um354, its modification level cannot inflate an OR calculation, and therefore, the Um354 edges provide saturation-independent evidence that the network is genuine. In 3D, the four residues reside within 25–35 Å of one another, and the cluster is distal to the decoding center, which is illustrated with the mRNA, A-, and P-site tRNAs (Figure 3i).

rRNA modifications are installed by two classes of guide snoRNAs, which are H/ACA box snoRNAs (SNORAs) for directing pseudouridylation (Ψ) by the isomerase dyskerin, and C/D box snoRNAs (SNORDs) for directing 2′-*O*-methylation (Nm) by the methyltransferase fibrillarin [46]. Next, we asked whether sharing a snoRNA guide could account for the coupling, since this would provide a clear mechanistic explanation for the elevated OR values. From the snoRNA database (snoDB) [47], 66 snoRNAs guide two or more rRNA modifications (Figure 4a).

**Figure 4.**
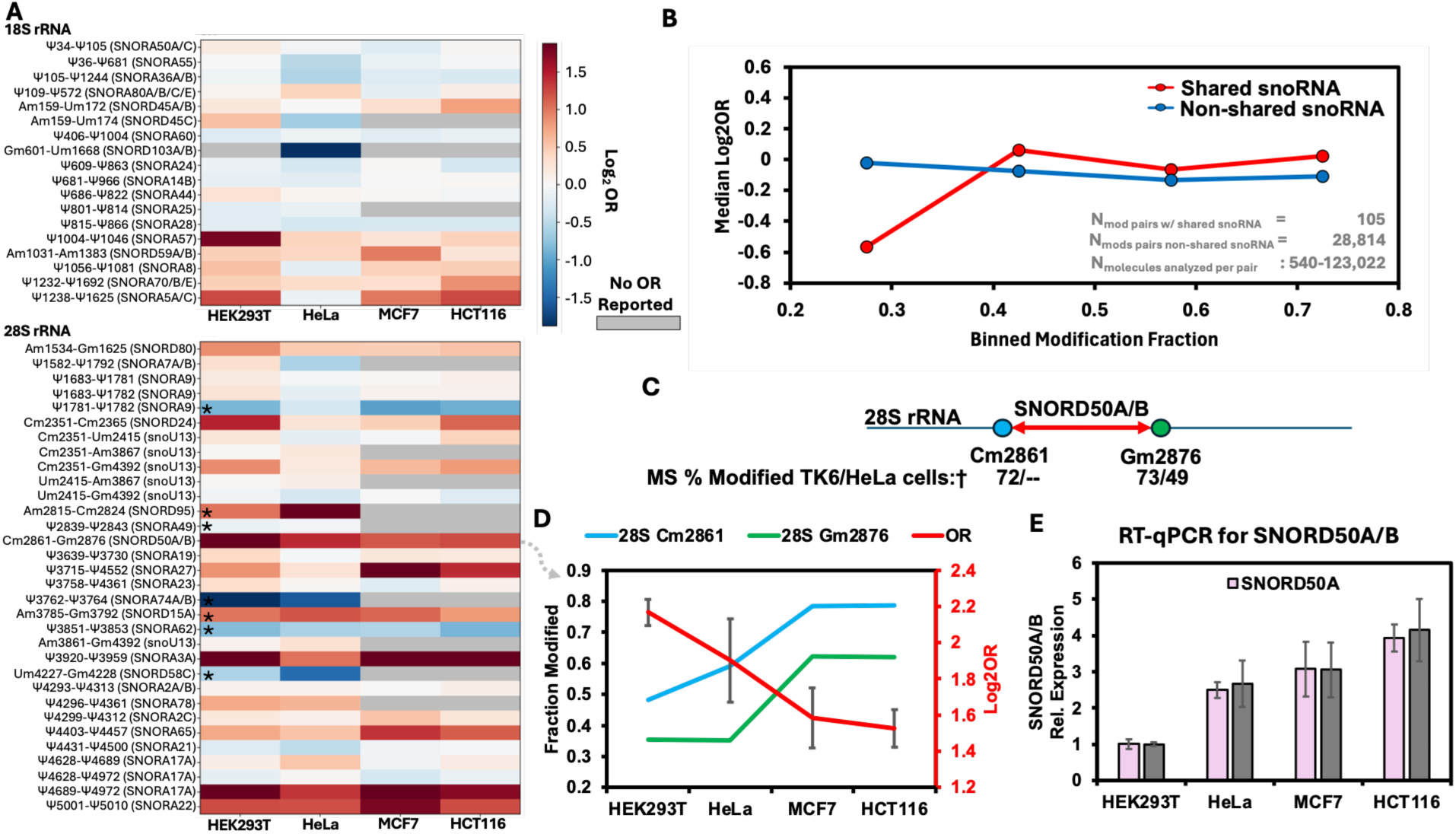
Ribosomal RNA modification co-occurrence is influenced by snoRNA guide availability in specific cases. (A) Heat map of log_2_OR values for each guide-sharing rRNA modification pair across the four cell lines. Asterisks mark pairs within 10 nt of one another, where k-mer effects can artifactually generate mutual exclusivity. (B) Median log2OR of modification pairs that share a snoRNA guide versus those that do not, binned by nanopore-measured modification fraction and pooled across the four cell lines. (C) Diagram of the 28S Cm2861–Gm2876 pair, both guided by SNORD50A/B, with the modification occupancies previously reported by mass spectrometry (†) [1]. (D) Nanopore-measured modification fractions for Cm2861 and Gm2876 and their log2OR values across the four cell lines. (E) Relative SNORD50A/B levels across the four cell lines measured by RT-qPCR.

The log_2_OR values for each snoRNA-sharing modification couple across the four cell lines were assembled into a heat map (Figure 4a). Sites marked with an asterisk are within 10 nt of one another in Figure 4a, which had intra-modification distances that can create k-mer effects that artificially generate mutual exclusivity (Figure 2i). The heat map does not reveal a consistent elevation in co-occurrence for modifications that share a snoRNA guide. To test this directly, we compared the median log_2_OR of guide-sharing pairs to that of pairs without a common guide, binned by modification fraction and pooled across the four cell lines (Figure 4b). Although the guide-sharing set of rRNA modifications is far smaller than the non-sharing set (105 versus 28,814 pairs), sharing a snoRNA guide did not increase the log_2_OR values for co-occurrence at matched modification fraction (Figure 4b). Thus, modifications sharing a snoRNA guide are indistinguishable from the background of other modification pairs.

We examined one guide-sharing pair in detail, which is the 28S rRNA Cm2861 and Gm2876 couple that are both methylated using the SNORD50A/B guides (Figure 4c). Across the four cell lines, the modification fractions from the Dorado Cm and Gm models revealed cell-line dependence at each site (Figure 4d). Cm2861 was least modified in HEK293T (0.48), intermediate in HeLa (0.59), and most modified in MCF7 and HCT116 (0.78). Gm2876 followed the same trend, low in HEK293T and HeLa (0.35 and 0.36) and higher in MCF7 and HCT116 (0.62). These nanopore values were consistent in direction with mass spectrometry reports for these sites, which found 28S 2861 to be 72% modified in TK6 cells and 28S 2876 to be 73% and 49% modified in TK6 and HeLa cells, respectively (Figure 4c) [1]. For Gm2876 in HeLa, the nanopore (35%) and mass spectrometry (49%; [1]) values differ by 14%, but both report a sub-stoichiometric level. All values are sub-stoichiometric and ideal for measuring co-occurrence. When the log_2_OR values were plotted across the four cell lines, co-occurrence was highest in HEK293T (2.16 ± 0.07), decreased in HeLa (1.9 ± 0.2), and was lowest in MCF7 (1.6 ± 0.2) and HCT116 (1.5 ± 0.1) cells (Figure 4d). Thus, co-occurrence and occupancy are inversely related across cell lines.

Next, whether the modification levels and their co-occurrence were related to the intracellular levels of SNORD50A/B was evaluated. Relative SNORD50A/B levels were measured by RT-qPCR against U6 snRNA as the internal standard (Figure 4e). SNORD50A/B was lowest in HEK293T cells, higher in HeLa cells, and maximal in MCF7 and HCT116 cells, which is the same rank order as the site occupancies and the inverse of the co-occurrence log_2_OR values. Thus, when the shared guide is limiting (HEK293T), both sites are modified at lower levels; however, they co-occur more often on the same molecule. In contrast, when the guide is abundant (MCF7, HCT116), the sites are more modified but with greater independence in the pool of strands. This pattern suggests that under limiting guide availability, the two modifications are written in an all-or-nothing manner on each rRNA molecule.

### Editing of rRNA modification networks with levofloxacin

A report from the Kool laboratory found that levofloxacin (Lev) binds a specific snoRNA in HEK293T cells, notably SNORD110 (Figure 5a) [48]. Binding to SNORD110 elevated its target modification, 18S Um1288. Following their example, HEK293T cells were exposed to 25 µM Lev for 48 h, followed by analysis of the rRNA with direct nanopore RNA sequencing to reveal whether the Lev-targeted modification changed and whether its co-occurrence with other modifications on individual molecules was altered.

**Figure 5.**
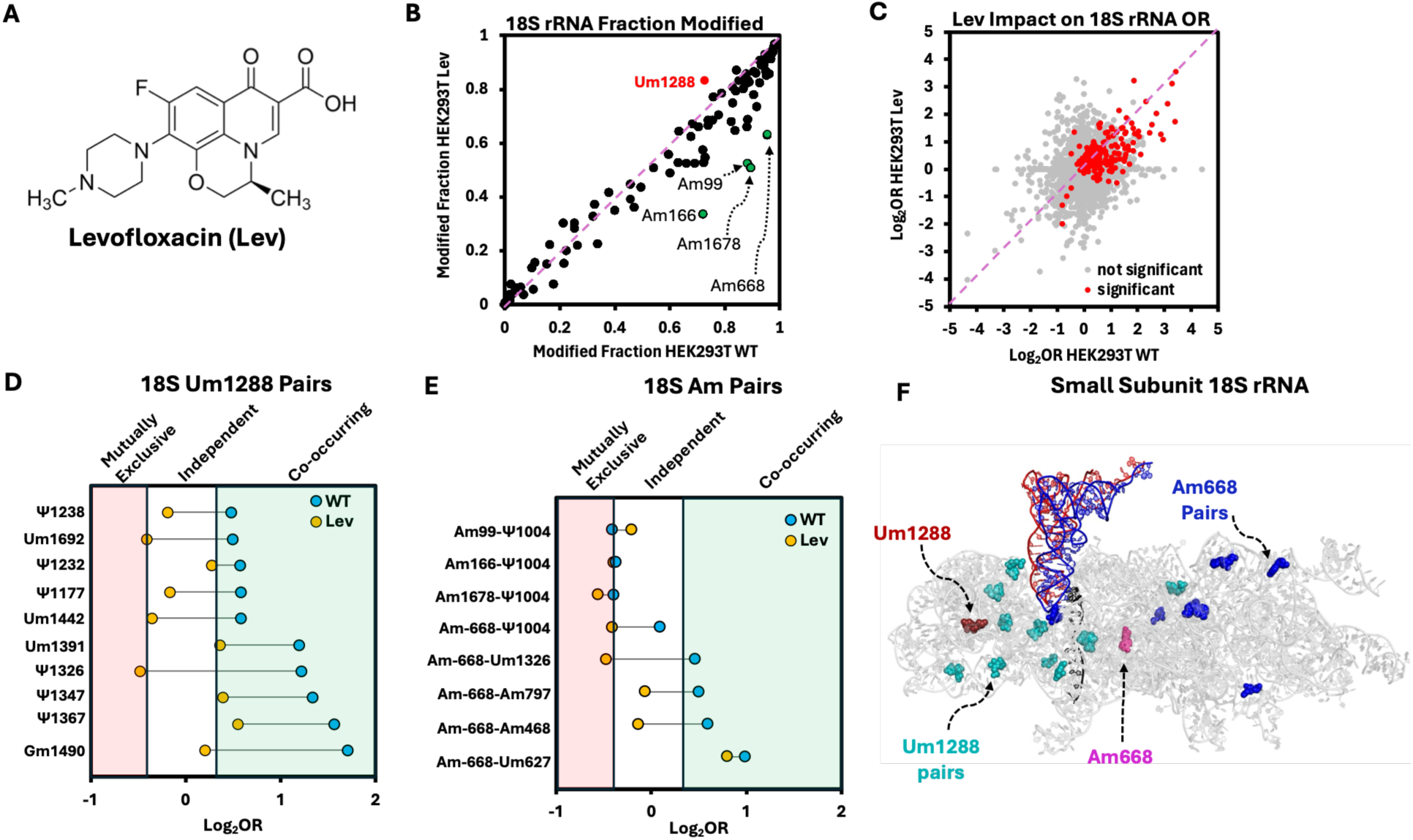
Treatment of HEK293T cells with levofloxacin alters rRNA modification levels and their co-occurrence. (A) Structure of levofloxacin. (B) Modification levels in the 18S rRNA from the wild-type versus Lev-treated HEK293T cells; red points mark the Um1288 modifications previously reported to change upon Lev treatment [48], and the circled points mark Am modifications found in the present work to change with Lev treatment. (C) Log2OR values in wild-type versus Lev-treated HEK293T cells for the 18S rRNA. The gray points are not significantly changed; the red points show statistically significant changes between conditions. (D) Forest plot of the co-occurrence changes for the 18S Um1288 network upon Lev treatment. (E) Forest plot for the 18S Am co-occurrence changes upon lev treatment. (F) rRNA sites impacted by lev treatment mapped onto the 18S rRNA structure (PDB 6Y0G) to visualize the sites are dispersed throughout the structure. The mRNA (black), A-(red), and P-site (blue) tRNAs are shown to provide a visual cue for the decoding center.

Comparison of the modification levels between wild-type and Lev-treated HEK293T cells found that U1288 increased by ∼10% when measured by nanopore analysis (Figure 5b), which is a smaller change than reported for the same cells and Lev treatment conditions when using a PCR-based assay of U1288 methylation (∼50%) [48]. It is not surprising that the two methods yield differences in the modification levels measured; nonetheless, both methods report the same direction of change.

Co-occurrence analysis revealed many co-occurring pairs in the 18S rRNA that decreased significantly upon Lev treatment (Figures 5c red points). A forest plot of the individual co-occurrence changes for Um1288, which is the known Lev-target methylation site, shows that co-occurrence with partner pairs decreased upon treatment. Seven of the ten members in the network were co-occurring before treatment and independent or mutually exclusive after Lev exposure.

In the modification fraction comparison, four 18S Am residues were found to decrease significantly in occupancy upon Lev treatment (positions 99, 166, 668, and 1678; Figure 5b). Inspection of these data to see whether co-occurrence occurs via these Am pairs on individual rRNA molecules identified Am668 as a hub for Am468, Um627, Cm797, and Ψ1004, as well as Ψ1004 forming co-occurring pairs with Am99, Am166, and Am1678 (Figure 5e). In general, after Lev treatment, the co-occurrence was independent or mutually exclusive compared to the wild-type state. The modifications identified were mapped onto the 18S rRNA structure to reveal they are dispersed throughout the small subunit ribosome and do not show a local favorability for co-occurrence as was identified in the previous analyses (Figure 3a). These data suggest a global impact of the drug on ribogenesis. In comparison, the SNORD50A/B levels across cell lines showed co-occurrence tracking with availability of the snoRNA guide (Figures 4d/e), while in the Lev-treated cells, the data support that perturbing the snoRNA machinery directly decoupled the target modifications from their networks, consistent with co-occurrence reflecting the availability of the modification machinery.

## Discussion

Ribosome biogenesis is among the most resource-intensive processes in the cell, requiring the coordinated action of all three RNA polymerases to assemble the rRNAs and proteins of the mature complex [49]. It is increasingly clear that the product of ribogenesis is not a single, uniform biomachine [6–10,14,15]. Human ribosomes vary in ribosomal-protein composition and their post-translational modification states, in rDNA-derived sequence variations in the rRNA pool, and in the levels of rRNA chemical modifications. Whether the rRNA modifications are installed independently or in coordinated patterns on individual rRNAs is not addressable by bulk measurements. Herein, modification-aware, single-molecule nanopore sequencing was used to measure rRNA modification co-occurrence one molecule at a time in human cytosolic rRNAs. To identify rRNA modification co-occurrence in the single-molecule data required controlling for the many known challenges of nanopore modification sequencing data analysis. An analytical framework was established that identifies co-occurrence by removing and controlling for read-quality effects, known false positives and calling artifacts, near-saturation that can masquerade as coupling, and global modification-level confounds (Figure 2).

The data analysis method was applied to the four human cell lines HEK293T, HeLa, MCF7, and HCT116. We found that co-occurrence is organized primarily by proximity, in that modifications close in sequence and 3D space tend to co-occur (Figure 3a). In contrast, modifications sharing a snoRNA guide are not preferentially written together on the same RNA strand compared to the population of modification pairs (Figures 4a/b). However, in one example (28S Cm2861-Gm2876 guided by SNORD50A/B), snoRNA guide availability and the rRNA modification levels for its targets show cell line-dependent results. Lastly, using a small molecule to perturb a target rRNA modification via its snoRNA, the single-molecule network involving that modification was diminished (Figures 4d/e). Together, these results establish per-molecule co-occurrence as a measurable layer of ribosome heterogeneity and begin to reveal how it is organized.

Global modification level possibly confounded the co-occurrence analysis and was removed by stratifying reads on their per-molecule modification level before computing the OR values (Figures 2 f/g). The distribution of these per-molecule levels is possibly interesting itself. The distribution of per-strand modification levels was ∼1.1-1.3x broader in the four cell lines than an analysis of 1,000 randomized samples of the data (Figures 2g and S6), which possibly indicates that the rRNA sites are modified with a degree of dependency. These data suggest there is more variation in rRNA modification levels between molecules than predicted by chance. The biological basis of this observation is not clear and was not further pursued because we cannot exclude contributions from all sequencing artifacts; moreover, rRNA sequence variations that exist in the data but were not interrogated introduce another level of heterogeneity in the rRNA that could impact the global distribution of per-strand modification levels.

Among the co-occurring pairs we identified (log_2_OR > 0.38; OR > 1.3), coupling was organized by spatial proximity. Pairs within ∼100 nucleotides of sequence and within ∼35–40 Å in 3D space were most likely to be written together (Figure 3a), suggesting that modifications are installed within a defined spatial window during ribogenesis and that a local region may mature as a unit. Importantly, coordination tracked with both sequence and 3D proximity, and its persistence within a ∼35–40 Å neighborhood argues that the relevant unit is a folded structural region rather than simply a stretch of primary sequence. This window is far smaller than the ∼600 Å that 100 nucleotides (0.6 Å per nucleotide) would span if fully extended, and it may correspond to a folding domain or an assembly intermediate accessible to a snoRNP. As a note, this value is empirically derived rather than a defined mechanistic length.

A key point is that the coupling observed is not an artifact of near saturation, as a ceiling threshold of 0.85 was applied to the data to avoid complications for which the mathematical limitation cannot be decoupled from biology (Figure 2e). Furthermore, 18S Um354 (∼20% modified by mass spectrometry and nanopore [1]) is an excellent example in which coupling was observed that changed with cell lines (Figures 3g/h). Finally, the mapped networks lie distal to the functional centers of the ribosome (Figures 3e/i), suggesting that an rRNA maturation intermediate [50] may lead to the molecule-to-molecule variation observed in the present work; nonetheless, peripheral modifications can still influence ribosome assembly, stability, and regulation [4,51]. Modifications known to reside near the peptidyl transferase and decoding centers exist at near-saturation levels and were not part of the analysis due to the ceiling threshold applied to the data; furthermore, at saturating levels, co-occurrence would not occur between molecules as they are all highly modified.

The co-occurrence networks were also cell-line dependent (Figures 3d/h), raising the possibility that the per-molecule modification patterns contribute to functional differences between ribosome populations; however, we did not test this hypothesis. The Um354 network is a useful example of the possibility of rRNA modification networks having functional consequences. Previous work from the Nova laboratory showed that in mouse cells the modification level of 18S Um355 (equivalent to human Um354) depends on developmental state and disease context of the cells studied [5]. Our data extend this observation with the finding that the modification fraction of 18S Um354 was human cell-line dependent, which was low in HEK293T and HCT116 cells and high in HeLa and MCF7 cells, and its co-occurrence with Ψ406, Um428, and Gm436 likewise differed between the cell lines (Figure 3g). Because this network spans both Nm and Ψ modifications and its three Nm sites are installed by different snoRNA guides (Ψ406 = SNORA60/71A-D; Um428 = SNIRD2A-C/68; Gm436 = SNORD100; and Um354 = SNORD90 [47]), its coordination again points to a spatial, rather than guide-based control over deposition of the modifications on rRNA molecules.

A striking feature of the data is that modification co-occurrence generally followed spatial proximity rather than shared snoRNA guides (Figures 3a and 4a/b), which might be assumed to couple their targets; however, there was one clear example of guide-dependent co-occurrence. The 28S modifications Cm2861 and Gm2876 are both methylated by fibrillarin using SNORD50A/B as the guide [47], and their co-occurrence was cell-line dependent with the coupling greatest in HEK293T cells and decreased through HeLa to MCF7 and HCT116 (Figures 4d/e). Using RT-qPCR to quantify SNORD50A/B levels, we found an inverse pattern between the snoRNA levels and Cm2861-Gm2876 co-occurrence (Figures 4d/e). This inverse relationship is consistent with prior work showing that SNORD50A/B (aka snoU50) levels are limiting for and directly set the methylation of Cm2861 (C2848 in the older numbering) [52]. Together, these observations indicate that, in the specific case of a limiting guide, the two modifications can be written in an all-or-nothing manner on each rRNA molecule.

Finally, we found that a small molecule binding a target snoRNA can not only alter a site-specific rRNA modification, consistent with the literature [48], but it can also change the coordination of the modification’s network across single molecules. Perturbing a specific snoRNA with Lev (SNORD110) produced a coherent decrease in the co-occurrence of the target modification with its network partners resulting in a state of independent installation of these modifications on rRNA molecules. This provides a complement to the SNORD50A/B observation, in which the coupling tracked guide availability across cell lines, whereas using Lev to perturb the snoRNA machinery directly reduced coupling. Unexpectedly, four Am residues (18S Am99, Am166, Am668, and Am1678; Figure 5b) were found to be modified at lower occupancy and decoupled co-occurrence after Lev treatment (Figure 5e), suggesting they are additional Lev-responsive sites. These sites do not share snoRNA guides (Am99 = SNORD57; Am166 = SNORD44; Am668 = SNORD36/36A-B; Am1678 = SNORD82 [47]); these data suggest Lev has a more global impact on rRNA modifications writing via altered writing kinetics, nucleolar stress, or stalled ribosome maturation. Regardless, Lev treatment leads to increased random writing of modifications on the final rRNA pool. Together with the prior studies, the data indicate that small molecules can alter an rRNA modification level via its snoRNA, while our data also identify the Lev-target modification’s single-molecule coordination is impacted.

Single-molecule nanopore sequencing has previously been applied to rRNA modification heterogeneity in yeast, using an HMM-based signal model to profile all modified positions per rRNA and pairwise rank correlations to identify concertedly modified positions [33]. The prior study and the present work both observe that rRNA modifications can be installed on individual molecules in a concerted manner, and the observations are only partly explained by a shared snoRNA guide. The prior and present analyses differ in organism (yeast versus human), sequencing chemistry, basecaller, and analytical framework (correlation-based profiling versus odds-ratio co-occurrence with explicit control for read-quality, saturation, and global modification-level confounds); therefore, further comparisons cannot be made.

These findings reframe rRNA modification not as a set of independent marks but as a partially coordinated feature of individual ribosomes, organized primarily by spatial proximity (Figure 3a), and in certain cases modulated by snoRNA guide availability (Figures 4d/e). An important caveat to these data is that co-occurrence is measured on mature cytoplasmic rRNA, so the spatial writing window inferred reflects the end state of modification rather than the act of installation; thus, the reported profiles reflect the pool of rRNAs that survived surveillance for incorrect synthesis, and the mature strands are the ones responsible for formation of specialized ribosomes. The finding of cell-line dependence in the co-occurrence of rRNA modifications on single RNA strands raises the possibility that modification neighborhoods contribute to a functionally heterogeneous ribosome pool unique to different cell types and states. The ability to read modification co-occurrence on single molecules, and to perturb them with a small molecule, provides a route to ask whether specific modification networks mark ribosome subpopulations and whether they can be manipulated for biological or therapeutic ends.

### Limitations

The Dorado base caller with the available modification-aware models was used, which has limitations, such as error-prone base calling and lack of complete modification-aware models [39,44,53]. Nonetheless, this tool is supported by ONT and easily implemented in high-performance computing centers for data analysis. We employed rigorous analysis protocols to minimize and remove the artifacts that arise in nanopore modification data, which include controlling for sequencing quality, per-molecule global modification level, local k-mer effects when more than one modification is read at once, and calling artifacts that produce false positives (Figures 2d-i). These address the known nanopore artifacts, but unknown artifacts may persist. A well-established issue not controlled for is that modification levels measured by nanopore sequencing do not always agree with those from mass spectrometry. For example, in HeLa cells, 18S Gm436 was 74% modified by nanopore but 59% by mass spectrometry, whereas 18S Um121 was 75% by nanopore but 98% by mass spectrometry (Figure S1; [1]); these examples show that the nanopore estimate can deviate in either direction. Other sites agree such as 18S Gm509 that was found to be 96% modified by both methods. Importantly, because these deviations are largely systematic, nanopore measurements are highly reproducible between replicates (Figures 2j/k), which is what enables reliable comparison between control and experimental samples. It is this reproducibility, rather than absolute accuracy, that underpins the analyses and conclusions reported here.

## Acknowledgements

The authors thank the University of Utah for startup funds that supported the project, and Prof. Cynthia Burrows at the University of Utah for many helpful discussions regarding the research.

## Supporting Information

Supporting information are available that include annotated rRNA modification sites analyzed, summary values for the analysis from all cell lines, plots of impact on data before and after quality and global confound analysis, global versus randomized sample analysis for each cell line, and 5.8S rRNA data analysis.

## Data Availability

The base-called nanopore data are publicly available at Zenodo doi:10.5281/zenodo.18602826 and doi:10.5281/zenodo.18602362. The code will be publicly deposited upon acceptance in a peer-reviewed journal.

## Conflict of Interests

The authors declare no competing financial interests in the reported studies.

## Notes

https://doi.org/10.5281/zenodo.18602826

https://doi.org/10.5281/zenodo.18602362

### Competing Interest Statement

The authors have declared no competing interest.

