## Supplemental Information for "Single-molecule nanopore sequencing reveals spatial coordination of rRNA modifications in human ribosomes"


| Item | Page |
| --- | --- |
| **Figure S1**. List of annotated rRNA epitranscriptomic modifications from the literature. | S2 |
| **Figure S2**. Summary data for the cell lines studied. | S6 |
| **Figure S3**. Summary values for the co-occurrence analysis. | S10 |
| **Figure S4**. HEK293T 28S rRNA data before and after data sequencing quality adjustment. | S11 |
| **Figure S5**. High occupancy and k-mer confounds in the 18S rRNA from HEK293T cells. | S12 |
| **Figure S6**. Distribution of global modification fraction versus a randomized sample. | S13 |
| **Figure S7.** Complete analysis of the 5.8S rRNA co-occurrence data set. | S18 |

**Figure S1**. List of annotated rRNA epitranscriptomic modifications from the literature.

| rRNA | Position | 28S Alt. Pos. | Modification | Taoka et al 2018 TK6 MS %Mod | Taoka et al 2018 HeLa MS % Mod | Holvec et al 2024 HeLa cryo-EM | Holm et al 2023 HEK293F cryo-EM | Faille et al 2023 HEK293F cryo-EM | Wiechert et al 2025 HeLa cryo-EM | Milenkovic et al 2025 Mouse nanopore unnanotated |
|---|---|---|---|---|---|---|---|---|---|---|
| 5.8S | 14 | | Um | 5 | 11 | | | | | |
| 5.8S | 55 | | Ψ | 60 | 80 | | | | | |
| 5.8S | 69 | | Ψ | 61 | 88 | | | | | |
| 5.8S | 75 | | Gm | 87 | 92 | | | | | |
| 18S | 27 | | Am | 100 | 96 | | | nd | nd | |
| 18S | 34 | | Ψ | 100 | 100 | | | nd | nd | |
| 18S | 36 | | Ψ | 82 | 93 | | | nd | nd | |
| 18S | 93 | | Ψ | 87 | 93 | | | nd | nd | |
| 18S | 99 | | Am | 99 | 94 | | | nd | nd | |
| 18S | 105 | | Ψ | 99 | 100 | | | nd | nd | |
| 18S | 109 | | Ψ | 99 | 99 | | | nd | nd | |
| 18S | 116 | | Um | 98 | 99 | | | nd | nd | |
| 18S | 119 | | Ψ | 94 | 98 | | | nd | nd | |
| 18S | 121 | | Um | 98 | 98 | | | nd | nd | |
| 18S | 159 | | Am | 96 | 92 | | | nd | nd | |
| 18S | 166 | | Am | 100 | 100 | | | nd | nd | |
| 18S | 172 | | Um | 96 | 100 | | | nd | nd | |
| 18S | 174 | | Cm | 92 | 91 | | | nd | nd | |
| 18S | 210 | | Ψ | 83 | 81 | | | | | |
| 18S | 218 | | Ψ | 100 | 100 | | | nd | nd | |
| 18S | 296 | | Ψ | 25 | nd | | | nd | nd | |
| 18S | 300 | | Ψ | nd | nd | | | nd | nd | |
| 18S | 354 | | Um | 20 | nd | | | nd | nd | |
| 18S | 406 | | Ψ | 87 | nd | | | nd | nd | |
| 18S | 428 | | Um | 76 | 78 | | | nd | nd | |
| 18S | 436 | | Gm | 76 | 57 | | | nd | nd | |
| 18S | 462 | | Cm | 100 | 100 | | | nd | nd | |
| 18S | 468 | | Am | 99 | 99 | | | nd | nd | |
| 18S | 484 | | Am | 97 | 100 | | | nd | nd | |
| 18S | 509 | | Gm | 98 | 96 | | | nd | nd | |
| 18S | 512 | | Am | 83 | nd | | | nd | nd | |
| 18S | 514 | | Ψ | nd | nd | | | nd | nd | |
| 18S | 517 | | Cm | 100 | 100 | | | nd | nd | |
| 18S | 556 | | Ψ | nd | nd | | | nd | nd | |
| 18S | 572 | | Ψ | 97 | 100 | | | nd | nd | |
| 18S | 573 | | Ψ | nd | nd | | | nd | nd | |
| 18S | 576 | | Am | 96 | 73 | | | nd | nd | |
| 18S | 581 | | U? | nd | nd | | | nd | nd | |
| 18S | 590 | | Am | 72 | 95 | | | nd | nd | |
| 18S | 601 | | Gm | 89 | 91 | | | nd | nd | |
| 18S | 609 | | Ψ | 90 | nd | | | nd | nd | |
| 18S | 621 | | Cm | 62 | nd | | | nd | nd | |
| 18S | 627 | | Um | 99 | nd | | | nd | nd | |
| 18S | 644 | | Gm | 98 | 100 | | | nd | nd | |
| 18S | 649 | | Ψ | 93 | 98 | | | nd | nd | |
| 18S | 651 | | Ψ | 93 | 98 | | | nd | nd | |
| 18S | 667 | | Ψ | nd | nd | | | nd | nd | |
| 18S | 668 | | Am | 99 | 100 | | | nd | nd | |
| 18S | 681 | | Ψ | 67 | 86 | | | nd | nd | |
| 18S | 683 | | Gm | 99 | nd | | | nd | nd | |
| 18S | 686 | | Ψ | 95 | 98 | | | nd | nd | |
| 18S | 769 | | Ψ | nd | nd | | | nd | nd | |
| 18S | 770 | | Ψ | nd | nd | | | nd | nd | |
| 18S | 797 | | Cm | 68 | nd | | | nd | nd | |
| 18S | 799 | | Um | 98 | 97 | | | nd | nd | |
| 18S | 801 | | Ψ | 100 | 100 | | | nd | nd | |
| 18S | 804 | | Ψ | nd | nd | | | nd | nd | |
| 18S | 814 | | Ψ | 100 | nd | | | nd | nd | |

Modification not Reported
Modification Reported
Unannotated Modification by Nanopore

| | | | | | | |
|---|---|---|---|---|---|---|
| 18S | 815 | Ψ | 100 | nd | nd | nd |
| 18S | 822 | Ψ | 99 | 99 | nd | nd |
| 18S | 863 | Ψ | 95 | 100 | nd | nd |
| 18S | 866 | Ψ | 88 | 97 | nd | nd |
| 18S | 867 | Gm | 28 | 48 | nd | nd |
| 18S | 889 | Ψ | nd | nd | nd | nd |
| 18S | 897 | Ψ | 23 | 33 | nd | nd |
| 18S | 918 | Ψ | 42 | nd | nd | nd |
| 18S | 966 | Ψ | 89 | 97 | nd | nd |
| 18S | 1003 | Ψ | nd | nd | nd | nd |
| 18S | 1004 | Ψ | 97 | 98 | nd | nd |
| 18S | 1031 | Am | 97 | 98 | nd | nd |
| 18S | 1045 | Ψ | 92 | nd | nd | nd |
| 18S | 1046 | Ψ | 100 | nd | nd | nd |
| 18S | 1056 | Ψ | 93 | 90 | nd | nd |
| 18S | 1061 | Ψ | nd | nd | nd | nd |
| 18S | 1081 | Ψ | 94 | 100 | nd | nd |
| 18S | 1136 | Ψ | 7 | <5 | nd | nd |
| 18S | 1174 | Ψ | 100 | 100 | nd | nd |
| 18S | 1177 | Ψ | 100 | nd | nd | nd |
| 18S | 1186 | Ψ | nd | nd | nd | nd |
| 18S | 1219 | C? | nd | nd | nd | nd |
| 18S | 1232 | Ψ | 98 | nd | nd | nd |
| 18S | 1238 | Ψ | 97 | 93 | nd | nd |
| 18S | 1239 | Ψ | nd | nd | nd | nd |
| 18S | 1244 | Ψ | 100 | 100 | nd | nd |
| 18S | 1248 | m1acp3Ψ | 100 | 100 | nd | nd |
| 18S | 1268 | C? | nd | nd | nd | nd |
| 18S | 1272 | Cm | 47 | nd | nd | nd |
| 18S | 1288 | Um | 98 | 72 | nd | nd |
| 18S | 1315 | Ψ | nd | nd | | |
| 18S | 1326 | Um | 100 | nd | nd | nd |
| 18S | 1328 | Gm | 100 | nd | nd | nd |
| 18S | 1337 | ac4C | 79 | nd | nd | nd |
| 18S | 1347 | Ψ | 98 | nd | nd | nd |
| 18S | 1359 | Ψ | nd | nd | nd | nd |
| 18S | 1367 | Ψ | 98 | 100 | nd | nd |
| 18S | 1383 | Am | 98 | nd | nd | nd |
| 18S | 1391 | Cm | 95 | 96 | nd | nd |
| 18S | 1400 | Ψ | nd | nd | nd | nd |
| 18S | 1442 | Um | 78 | 96 | nd | nd |
| 18S | 1445 | Ψ | 90 | 100 | nd | nd |
| 18S | 1447 | Gm | 34 | 39 | nd | nd |
| 18S | 1463 | U? | nd | nd | nd | nd |
| 18S | 1490 | Gm | 100 | 100 | nd | nd |
| 18S | 1596 | Ψ | nd | nd | nd | nd |
| 18S | 1625 | Ψ | 79 | 69 | nd | nd |
| 18S | 1639 | m7G | 100 | 100 | nd | nd |
| 18S | 1643 | Ψ | 96 | 92 | nd | nd |
| 18S | 1668 | Um | 8 | <5 | nd | nd |
| 18S | 1678 | Am | 94 | nd | nd | nd |
| 18S | 1692 | Ψ | 98 | 100 | nd | nd |
| 18S | 1703 | Cm | 92 | nd | nd | nd |
| 18S | 1804 | Um | 86 | 80 | nd | nd |
| 18S | 1832 | m6A | 99 | 100 | nd | nd |
| 18S | 1839 | Ψ | nd | nd | nd | nd |
| 18S | 1842 | ac4C | 99 | 97 | nd | nd |
| 18S | 1850 | m62A | 94 | 99 | nd | nd |
| 18S | 1851 | m62A | 94 | 99 | nd | nd |

| | | | | Taoka et al 2018 | | Holvec et al 2024 | Holm et al 2023 | Faille et al 2023 | Wiechert et al 2025 | Milenkovic et al 2025 |
|---|---|---|---|---|---|---|---|---|---|---|
| rRNA | Position | 28S Alt. Pos. | Modification | TK6 MS %Mod | HeLa MS % Mod | HeLa cryo-EM | HEK293F cryo-EM | HEK293F cryo-EM | HeLa cryo-EM | Mouse nanopore unnanotated |
| 28S | 224 | | Ψ | nd | nd | | | | | |
| 28S | 398 | 389 | Am | 98 | 97 | | | | | |
| 28S | 400 | 391 | Am | 98 | 100 | | | | | |
| 28S | 1316 | 1303 | Gm | 71 | 36 | | | | | |
| 28S | 1322 | 1309 | m1A | 100 | 100 | | | | | |
| 28S | 1323 | 1310 | Am | 44 | 27 | | | | | |
| 28S | 1326 | 1313 | Am | 100 | 100 | | | | | |
| 28S | 1340 | 1327 | Cm | 92 | 87 | | | | | |
| 28S | 1522 | 1509 | Gm | 99 | 99 | | | | | |
| 28S | 1524 | 1511 | Am | 99 | 98 | | | | | |
| 28S | 1534 | 1521 | Am | 100 | 100 | | | | | |
| 28S | 1536 | 1523 | Ψ | 88 | 78 | | | | | |
| 28S | 1582 | 1569 | Ψ | 68 | nd | | | | | |
| 28S | 1625 | 1612 | Gm | 100 | 100 | | | | | |
| 28S | 1677 | 1664 | Ψ | 97 | 98 | | | | | |
| 28S | 1683 | 1670 | Ψ | 96 | 93 | | | | | |
| 28S | 1700 | 1687 | U? | nd | nd | | | | | |
| 28S | 1744 | 1731 | Ψ | 100 | nd | | | | | |
| 28S | 1760 | 1747 | Gm | 89 | nd | | | | | |
| 28S | 1773 | 1760 | Um | 70 | nd | | | | | |
| 28S | 1779 | 1766 | Ψ | 40 | nd | | | | | |
| 28S | 1781 | 1768 | Ψ | 100 | nd | | | | | |
| 28S | 1782 | 1769 | Ψ | 100 | nd | | | | | |
| 28S | 1792 | 1779 | Ψ | 100 | nd | | | | | |
| 28S | 1859 | 1846 | m4C | | | | | | | |
| 28S | 1860 | 1847 | Ψ | 95 | 98 | | | | | |
| 28S | 1862 | 1849 | Ψ | 95 | 94 | | | | | |
| 28S | 1871 | 1858 | Am | 96 | 94 | | | | | |
| 28S | 1881 | 1868 | Cm | 35 | nd | | | | | |
| 28S | 2351 | 2338 | Cm | 99 | nd | | | | | |
| 28S | 2363 | 2350 | Am | 100 | 100 | | | | | |
| 28S | 2364 | 2351 | Gm | 100 | 100 | | | | | |
| 28S | 2365 | 2352 | Cm | 90 | 96 | | | | | |
| 28S | 2401 | 2388 | Am | 73 | 66 | | | | | |
| 28S | 2415 | 2402 | Um | 87 | nd | | | | | |
| 28S | 2422 | 2409 | Cm | 98 | nd | | | | | |
| 28S | 2424 | 2411 | Gm | 90 | nd | | | | | |
| 28S | 2508 | 2495 | Ψ | 92 | 76 | | | | | |
| 28S | 2632 | 2619 | Ψ | 96 | 93 | | | | | |
| 28S | 2787 | 2774 | Am | 84 | 78 | | | | | |
| 28S | 2804 | 2791 | Cm | 93 | 89 | | | | | |
| 28S | 2815 | 2802 | Am | 92 | 78 | | | | | |
| 28S | 2824 | 2811 | Cm | 87 | 66 | | | | | |
| 28S | 2837 | 2824 | Um | 99 | 99 | | | | | |
| 28S | 2839 | 2826 | Ψ | 20 | nd | | | | | |
| 28S | 2843 | 2830 | Ψ | 9 | <5 | | | | | |
| 28S | 2856 | 2843 | U? | nd | nd | | | | | |
| 28S | 2861 | 2848 | Cm | 72 | nd | | | | | |
| 28S | 2876 | 2863 | Gm | 49 | 73 | | | | | |
| 28S | 3627 | 3606 | Gm | 96 | 84 | | | | | |
| 28S | 3637 | 3616 | Ψ | 89 | nd | | | | | |
| 28S | 3639 | 3618 | Ψ | 95 | 96 | | | | | |
| 28S | 3669 | 3648 | Gm | nd | nd | | | | | |
| 28S | 3695 | 3674 | Ψ | 99 | 100 | | | | | |
| 28S | 3701 | 3680 | Cm | 100 | nd | | | | | |
| 28S | 3715 | 3694 | Ψ | 100 | nd | | | | | |
| 28S | 3718 | 3697 | Am | 88 | 87 | | | | | |
| 28S | 3723 | 3702 | Am | 100 | 100 | | | | | |
| 28S | 3724 | 3703 | Am | nd | nd | | | | | |
| 28S | 3730 | 3709 | Ψ | 72 | nd | | | | | |
| 28S | 3734 | 3713 | Ψ | 98 | nd | | | | | |
| 28S | 3744 | 3723 | Gm | 83 | 94 | | | | | |
| 28S | 3758 | 3737 | Ψ | 85 | 100 | | | | | |
| 28S | 3760 | 3739 | Am | 90 | 100 | | | | | |
| 28S | 3762 | 3741 | Ψ | 100 | 100 | | | | | |
| 28S | 3764 | 3743 | Ψ | 100 | 100 | | | | | |
| 28S | 3768 | 3747 | Ψ | 100 | 100 | | | | | |
| 28S | 3770 | 3749 | Ψ | 100 | 100 | | | | | |

| | | | | | |
|---|---|---|---|---|---|
| 28S | 3782 | 3761 | m5C | 100 | 100 |
| 28S | 3785 | 3764 | Am | 96 | 100 |
| 28S | 3792 | 3771 | Gm | 100 | 100 |
| 28S | 3808 | 3787 | Cm | 80 | nd |
| 28S | 3818 | 3797 | Ψm | 100 | nd |
| 28S | 3822 | 3801 | Ψ | 50 | nd |
| 28S | 3825 | 3804 | Am | 92 | nd |
| 28S | 3830 | 3809 | Am | 100 | 98 |
| 28S | 3841 | 3820 | Cm | 100 | 100 |
| 28S | 3844 | 3823 | Ψ | 66 | 100 |
| 28S | 3851 | 3830 | Ψ | 92 | 100 |
| 28S | 3853 | 3832 | Ψ | 100 | 100 |
| 28S | 3867 | 3846 | Am | 43 | 40 |
| 28S | 3869 | 3848 | Cm | 67 | 93 |
| 28S | 3884 | 3863 | Ψ | 33 | nd |
| 28S | 3887 | 3866 | Cm | 99 | 100 |
| 28S | 3899 | 3878 | Gm | 98 | 98 |
| 28S | 3920 | 3899 | Ψ | 100 | nd |
| 28S | 3925 | 3904 | Um | 96 | 92 |
| 28S | 3944 | 3923 | Gm | 80 | 77 |
| 28S | 3959 | 3938 | Ψ | 93 | nd |
| 28S | 4042 | 4020 | Gm | 83 | nd |
| 28S | 4054 | 4032 | Cm | 100 | 100 |
| 28S | 4196 | 4166 | Gm | 98 | nd |
| 28S | 4220 | 4190 | m6A | 100 | 100 |
| 28S | 4227 | 4197 | Um | 97 | nd |
| 28S | 4228 | 4198 | Gm | 92 | nd |
| 28S | 4293 | 4263 | Ψ | 98 | 100 |
| 28S | 4296 | 4266 | Ψ | 90 | 90 |
| 28S | 4299 | 4269 | Ψ | 93 | 100 |
| 28S | 4306 | 4276 | Um | 88 | nd |
| 28S | 4312 | 4282 | Ψ | 83 | 94 |
| 28S | 4353 | 4323 | Ψ | 95 | 99 |
| 28S | 4361 | 4331 | Ψ | 93 | 82 |
| 28S | 4370 | 4340 | Gm | 99 | nd |
| 28S | 4392 | 4362 | Gm | 97 | 96 |
| 28S | 4403 | 4373 | Ψ | 96 | nd |
| 28S | 4420 | 4390 | Ψ | 99 | 94 |
| 28S | 4423 | 4393 | Ψ | 97 | 100 |
| 28S | 4431 | 4401 | Ψ | 89 | nd |
| 28S | 4442 | 4412 | Ψ | 100 | 100 |
| 28S | 4447 | 4417 | m5C | 100 | 100 |
| 28S | 4456 | 4426 | Cm | 98 | 97 |
| 28S | 4457 | 4427 | Ψ | 98 | 100 |
| 28S | 4471 | 4441 | Ψ | 87 | nd |
| 28S | 4493 | 4463 | Ψ | 17 | nd |
| 28S | 4494 | 4464 | Gm | 91 | nd |
| 28S | 4498 | 4468 | Um | 100 | 100 |
| 28S | 4499 | 4469 | Gm | 100 | 100 |
| 28S | 4500 | 4470 | Ψ | 100 | 100 |
| 28S | 4521 | 4491 | Ψ | 91 | nd |
| 28S | 4523 | 4493 | Am | 87 | nd |
| 28S | 4530 | 4500 | m3U | 100 | 100 |
| 28S | 4531 | 4501 | Ψ | nd | nd |
| 28S | 4532 | 4502 | Ψ | 100 | 100 |
| 28S | 4536 | 4506 | Cm | 100 | nd |
| 28S | 4552 | 4522 | Ψ | 98 | 92 |
| 28S | 4569 | 4539 | Ψ | nd | nd |
| 28S | 4571 | 4541 | Am | 43 | nd |
| 28S | 4576 | 4546 | Ψ | 100 | nd |
| 28S | 4579 | 4549 | Ψ | 100 | nd |
| 28S | 4590 | 4560 | Am | 37 | 32 |
| 28S | 4599 | 4569 | U? | nd | nd |
| 28S | 4618 | 4588 | Gm | 75 | nd |
| 28S | 4620 | 4590 | Um | 82 | nd |
| 28S | 4623 | 4593 | Gm | 100 | nd |
| 28S | 4628 | 4598 | Ψ | 92 | nd |
| 28S | 4636 | 4606 | Ψ | 42 | nd |
| 28S | 4637 | 4607 | Gm | 100 | nd |
| 28S | 4673 | 4643 | Ψ | 39 | 36 |
| 28S | 4689 | 4659 | Ψ | 87 | nd |
| 28S | 4972 | 4937 | Ψ | 81 | 92 |
| 28S | 4973 | 4938 | Ψ | nd | nd |
| 28S | 5001 | 4966 | Ψ | 86 | nd |
| 28S | 5010 | 4975 | Ψ | 72 | nd |

**Figure S2**. Summary data for the cell lines studied.

MCF7-1

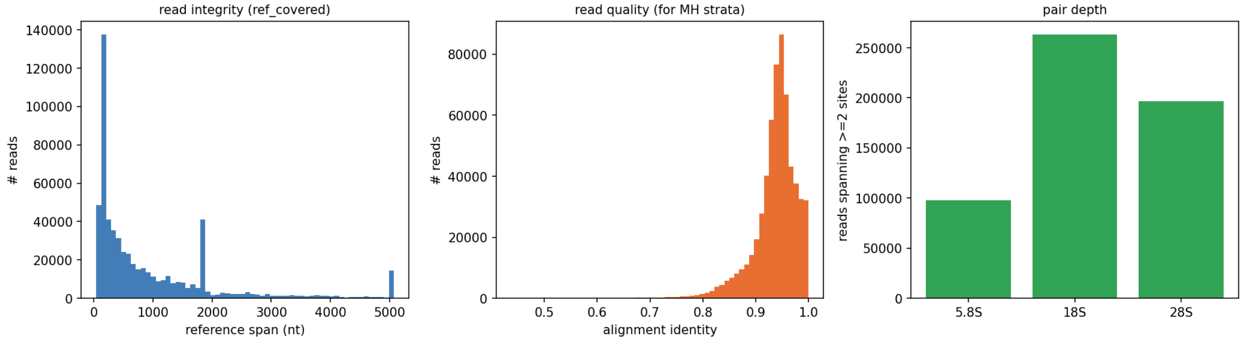


MCF7-2

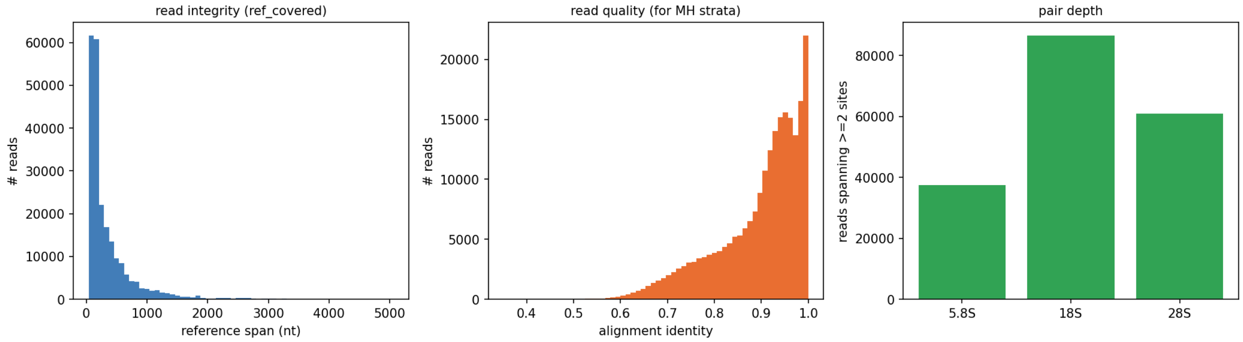


HEK293T-1

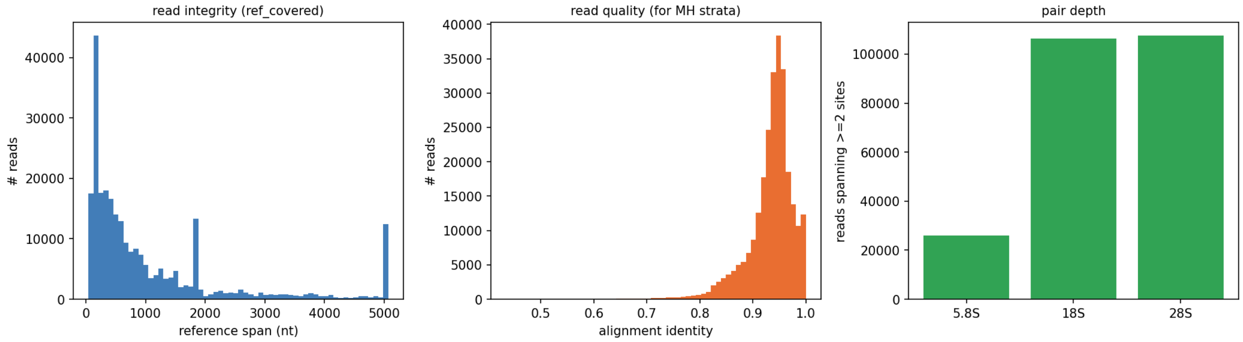


HEK293T-2

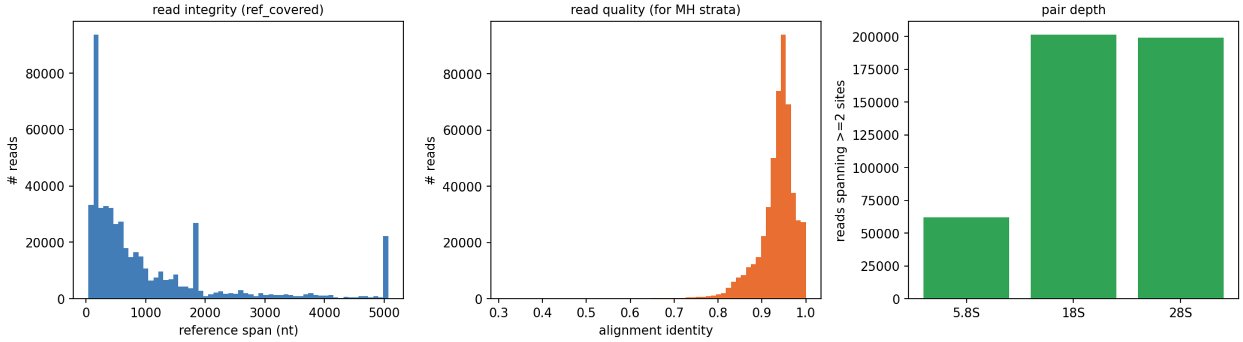


HeLa-1

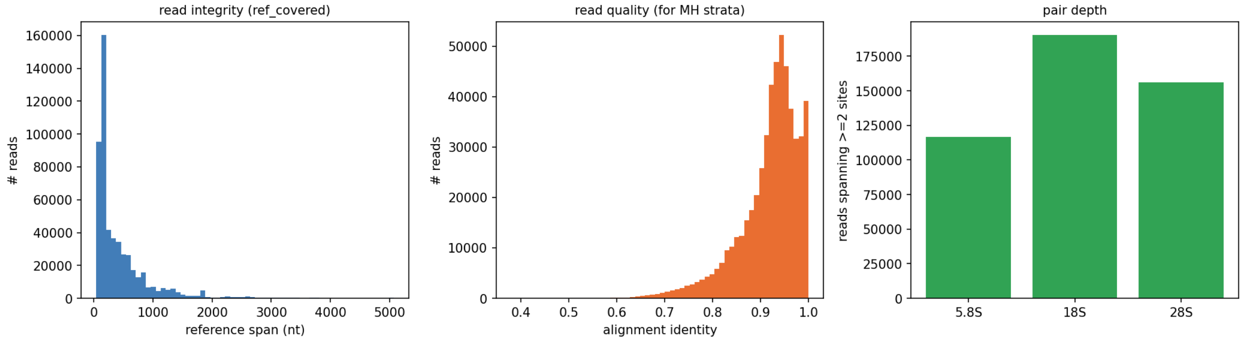


HeLa-2

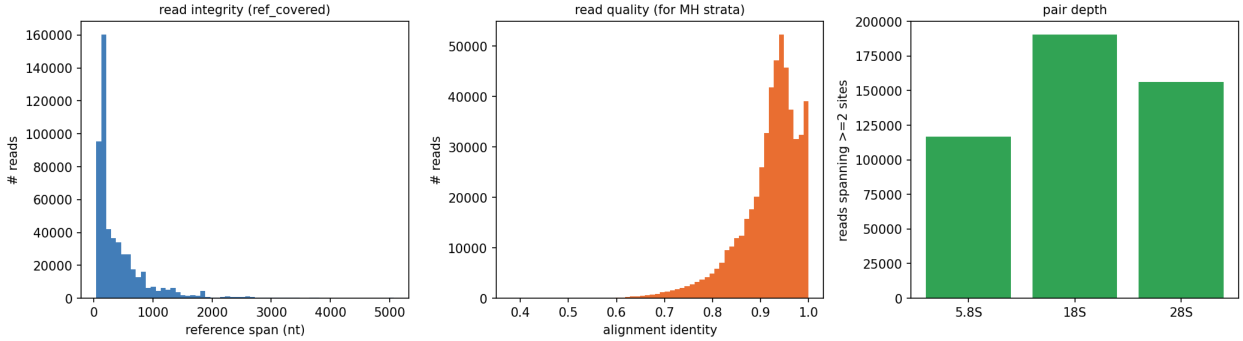


HCT116-1

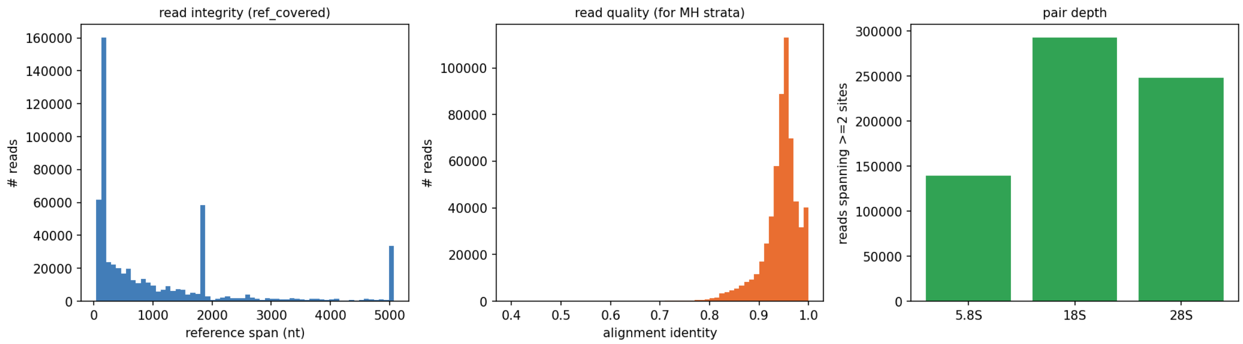


HCT116-2

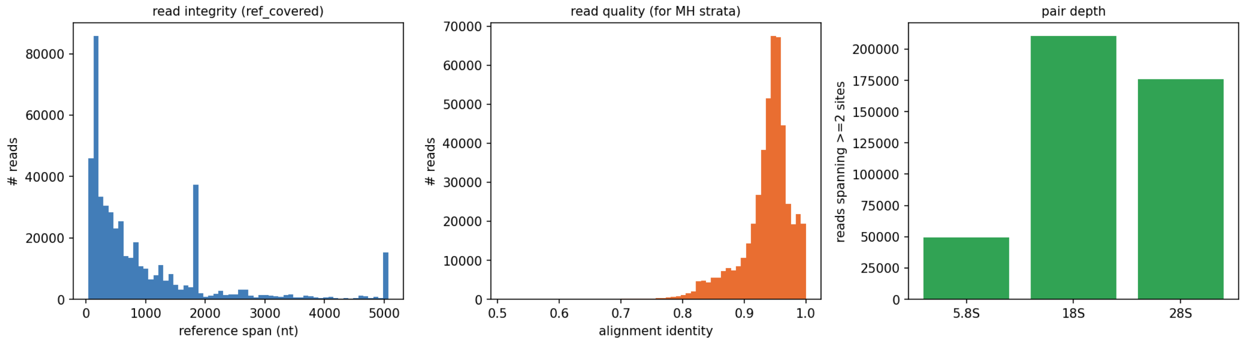


HEK293T-Lev-1

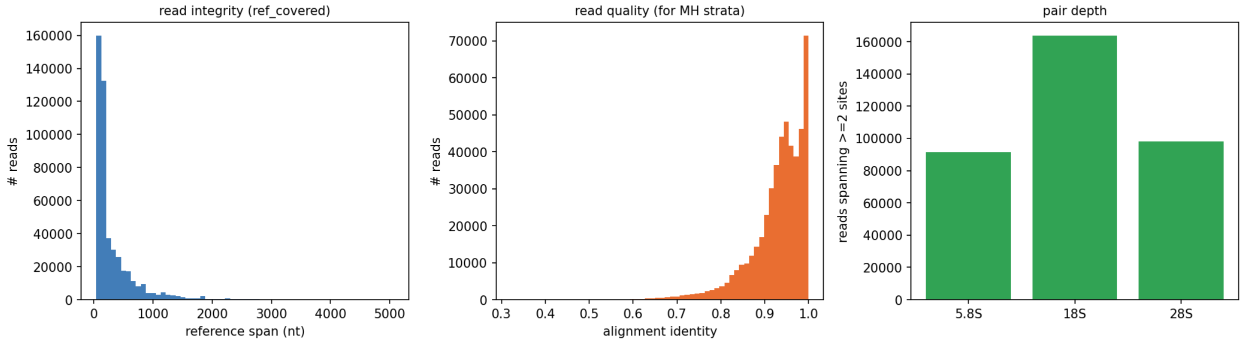


HEK293T-Lev-2

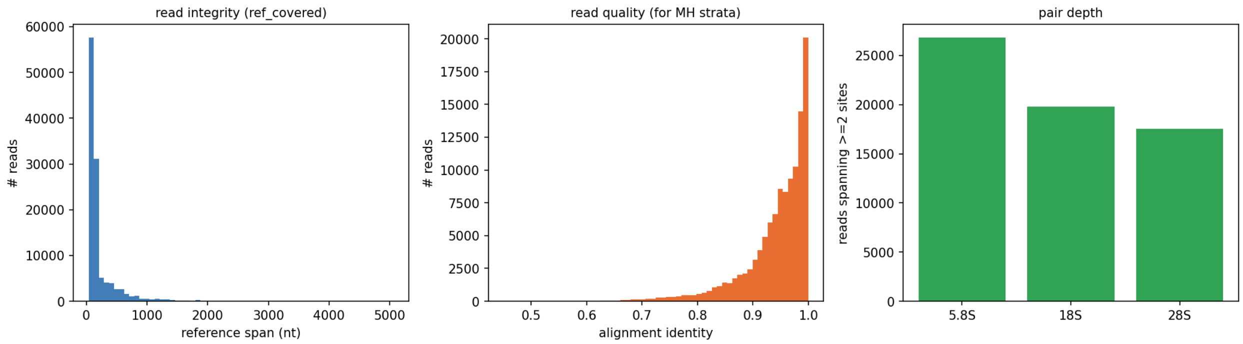
read integrity (ref_covered)
### reads
60000
50000
40000
30000
20000
10000
0
0
1000
2000
3000
4000
5000
reference span (nt)
read quality (for MH strata)
### reads
20000
17500
15000
12500
10000
7500
5000
2500
0
0.5
0.6
0.7
0.8
0.9
1.0
alignment identity
pair depth
reads spanning >=2 sites
25000
20000
15000
10000
5000
0
5.8S
18S
28S

**Figure S3**. Summary values for the co-occurrence analysis.

| strand | cell_line | n_pairs_total | n_within_kmer | n_lowvar_pairs | n_global_confounded | n_OR_nonfinite | n_analyzable | n_cooc_signif. | %_cooc_signif. | depth_min | depth_median | depth_max |
|---|---|---|---|---|---|---|---|---|---|---|---|---|
| **18S** | HEK293T-1 | 5356 | 131 | 0 | 150 | 63 | 5021 | 226 | 4.5 | 1681 | 13156 | 67146 |
| **18S** | HEK293T-2 | 5356 | 131 | 0 | 138 | 24 | 5070 | 276 | 5.44 | 3405 | 26457 | 137335 |
| **18S** | HEK293T-Lev-1 | 5356 | 131 | 0 | 91 | 97 | 5048 | 54 | 1.07 | 169 | 2795.5 | 76307 |
| **18S** | HEK293T-Lev-2 | 5356 | 131 | 0 | 6 | 820 | 4415 | 6 | 0.14 | 16 | 330 | 8868 |
| **18S** | HeLa-1 | 5356 | 131 | 0 | 56 | 59 | 5118 | 132 | 2.58 | 215 | 5450.5 | 87565 |
| **18S** | HeLa-2 | 5356 | 131 | 0 | 73 | 54 | 5106 | 133 | 2.6 | 185 | 5371.5 | 87607 |
| **18S** | MCF7-1 | 2628 | 68 | 675 | 74 | 0 | 1857 | 120 | 6.46 | 3289 | 34262.5 | 123022 |
| **18S** | MCF7-2 | 5356 | 131 | 0 | 53 | 0 | 4947 | 260 | 5.25 | 18 | 1057.5 | 29618 |
| **18S** | HCT116-1 | 2628 | 68 | 675 | 161 | 0 | 1808 | 70 | 3.87 | 3078 | 23737 | 109724 |
| **18S** | HCT116-2 | 2628 | 68 | 675 | 335 | 0 | 1688 | 97 | 5.75 | 3497 | 31080.5 | 149812 |
| **28S** | HEK293T-1 | 9045 | 212 | 0 | 913 | 4 | 7938 | 627 | 7.9 | 482 | 16874 | 64920 |
| **28S** | HEK293T-2 | 9045 | 212 | 0 | 1071 | 0 | 7780 | 796 | 10.23 | 773 | 30656 | 124171 |
| **28S** | HEK293T-Lev-1 | 9045 | 212 | 0 | 609 | 340 | 7928 | 50 | 0.63 | 7 | 1675 | 36259 |
| **28S** | HEK293T-Lev-2 | 9045 | 212 | 0 | 31 | 1746 | 7077 | 11 | 0.16 | 0 | 149 | 5618 |
| **28S** | HeLa-1 | 9045 | 212 | 0 | 266 | 169 | 8423 | 149 | 1.77 | 16 | 3353 | 63154 |
| **28S** | HeLa-2 | 9045 | 212 | 0 | 399 | 168 | 8290 | 178 | 2.15 | 21 | 3338 | 63412 |
| **28S** | MCF7-1 | 9045 | 121 | 693 | 875 | 0 | 7531 | 653 | 8.67 | 517 | 26401 | 97928 |
| **28S** | MCF7-2 | 9045 | 212 | 0 | 204 | 0 | 8109 | 601 | 7.41 | 2 | 527 | 16699 |
| **28S** | HCT116-1 | 5253 | 121 | 693 | 660 | 0 | 3955 | 365 | 9.23 | 751 | 21604 | 89297 |
| **28S** | HCT116-2 | 5253 | 121 | 693 | 900 | 0 | 3780 | 343 | 9.07 | 857 | 25062 | 104795 |

**Figure S4**. HEK293T 28S rRNA data before and after data sequencing quality adjustment.

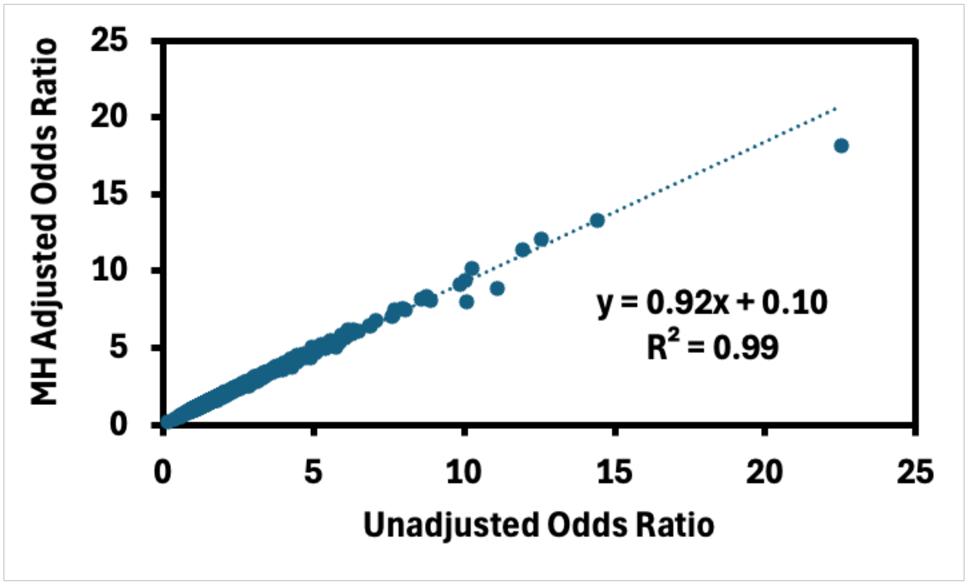

**Figure S5**. High occupancy and k-mer confounds in the 18S rRNA from HEK293T cells.

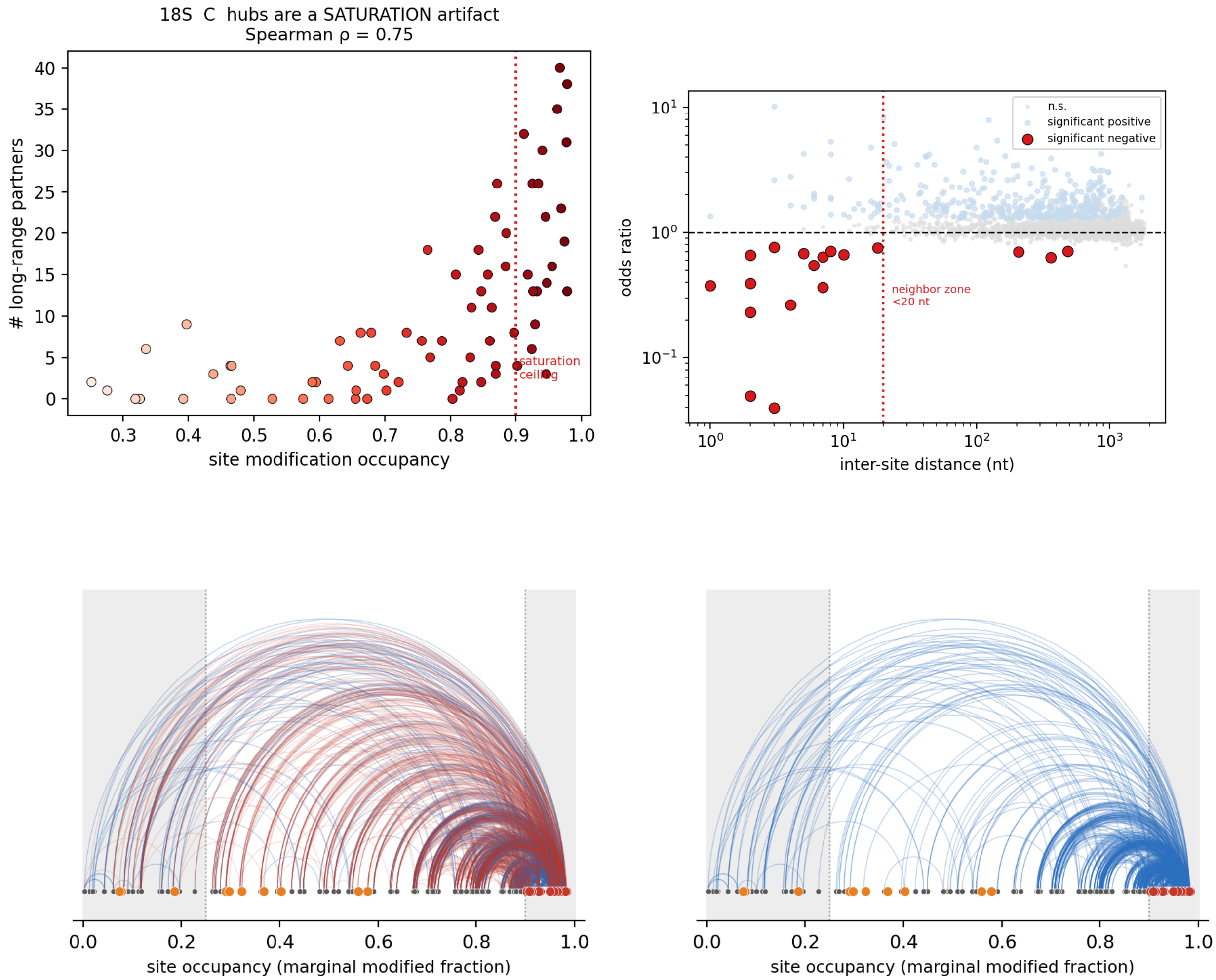


**Example 28S funnel (positive co-occurrence):**

| filter | pairs remaining | removed |
|---|---|---|
| all pairs | 9,045 | |
| positive (OR ≥ 1) | 7,012 | |
| **+ survive OR_adj** (global survivors) | **841** | |
| + both sites sub-saturated (0.25–0.85) | **13** | 828 |
| + non-k-mer (dist > 5) | 9 | 4 |

**Figure S6**. Distribution of global modification fraction versus a randomized sample.

HEK293T 18S rRNA

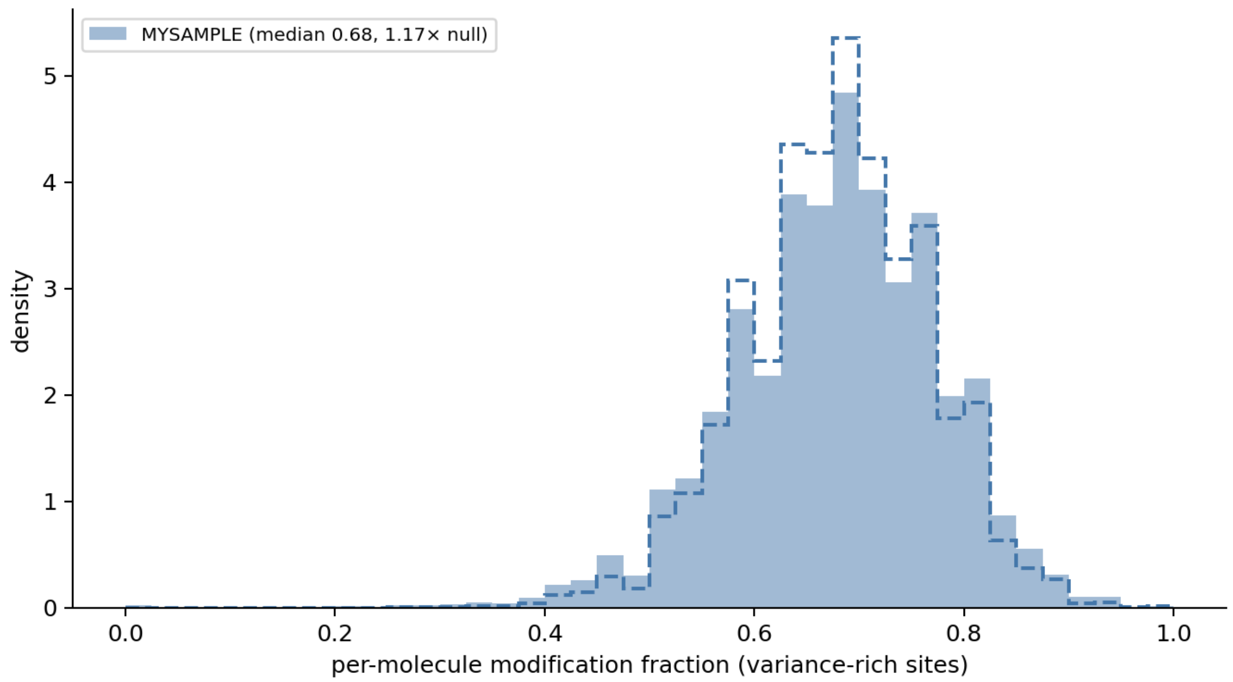


HEK293T 28S rRNA

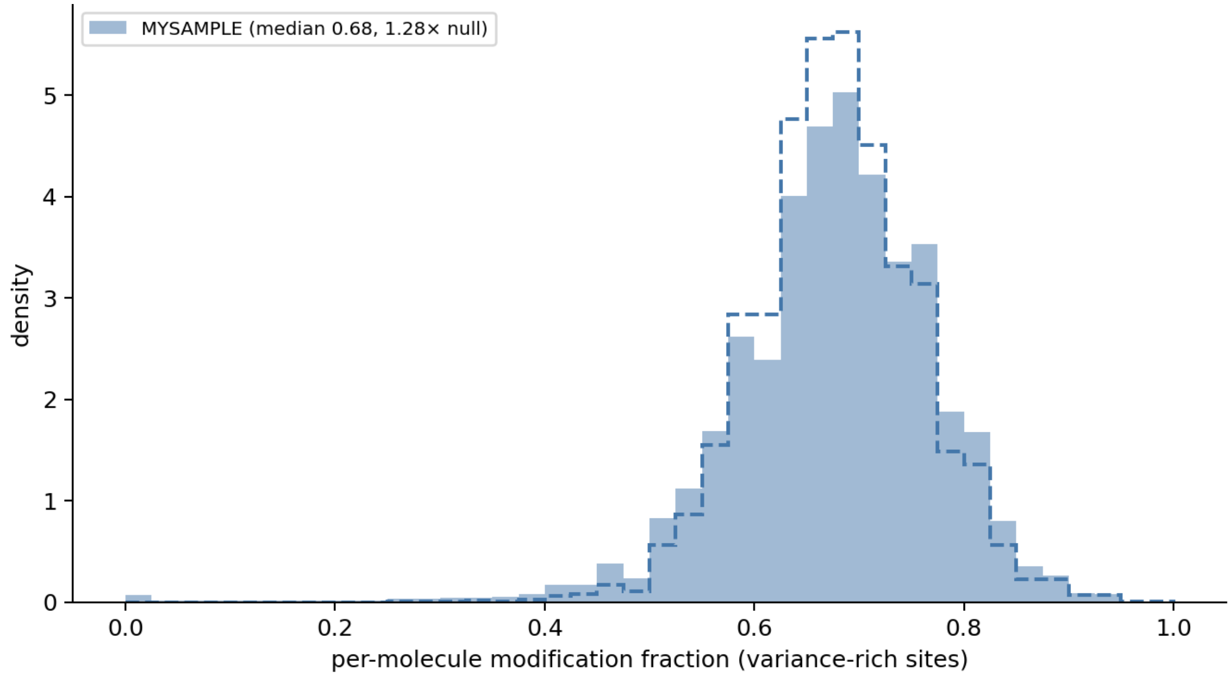


MCF7 18S rRNA

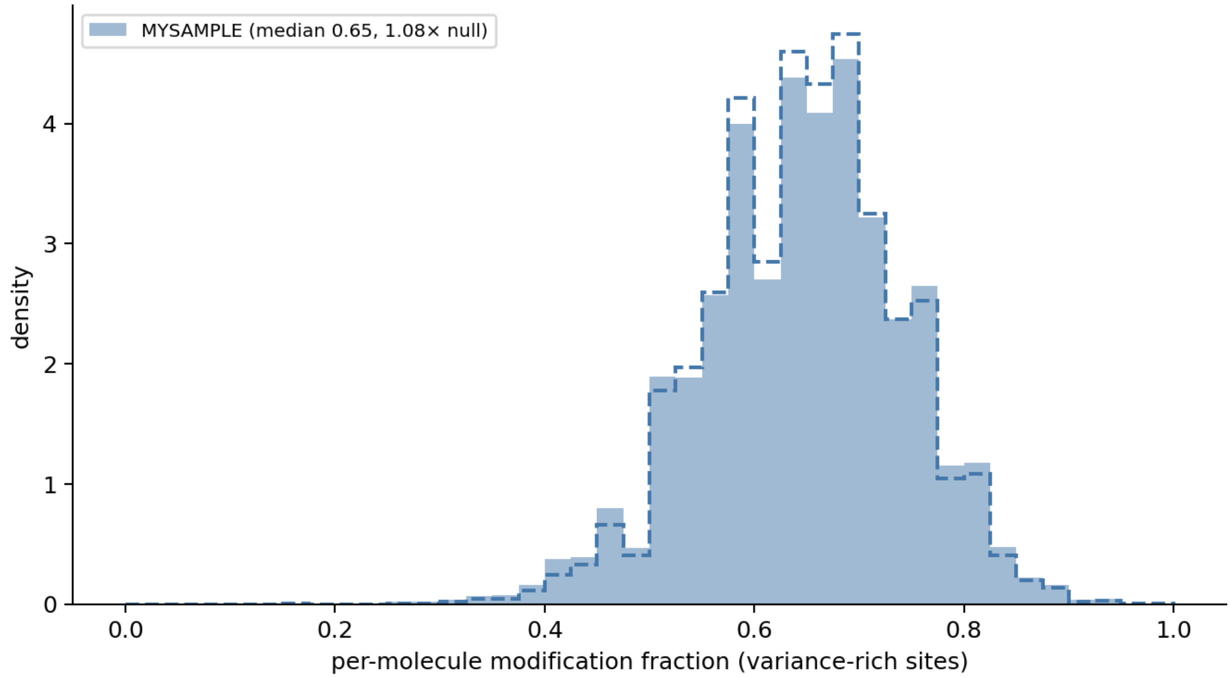


MCF7 28S rRNA

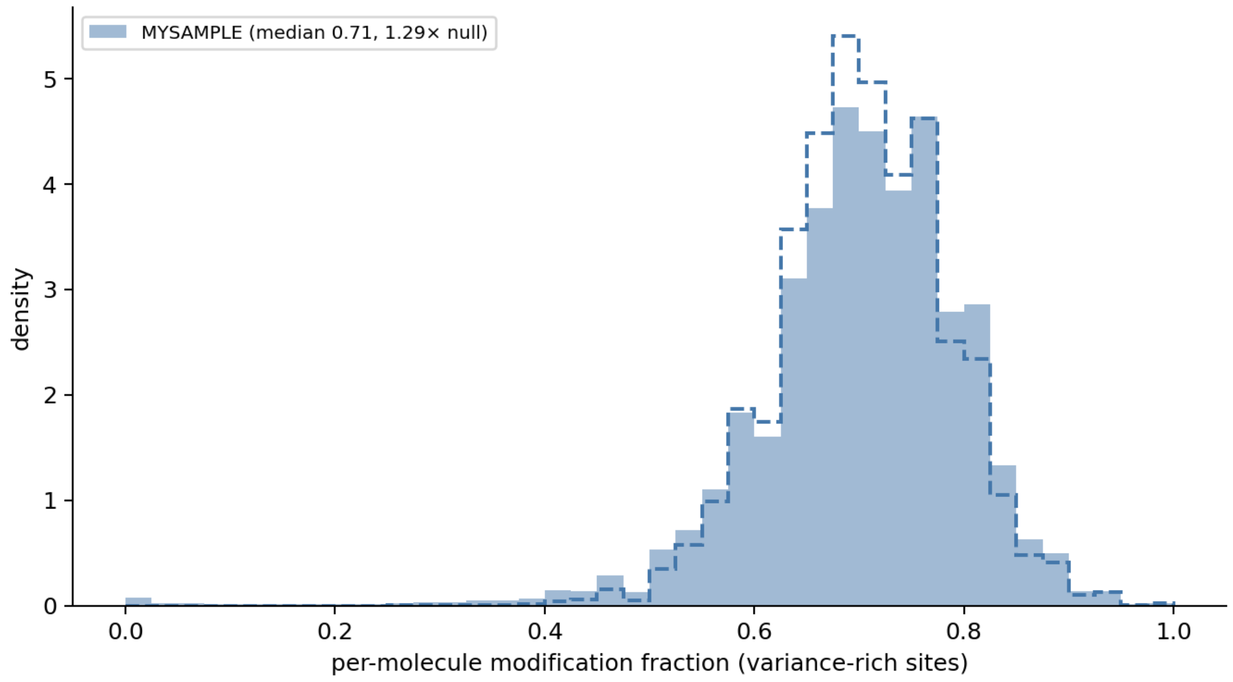


HeLa 18S rRNA

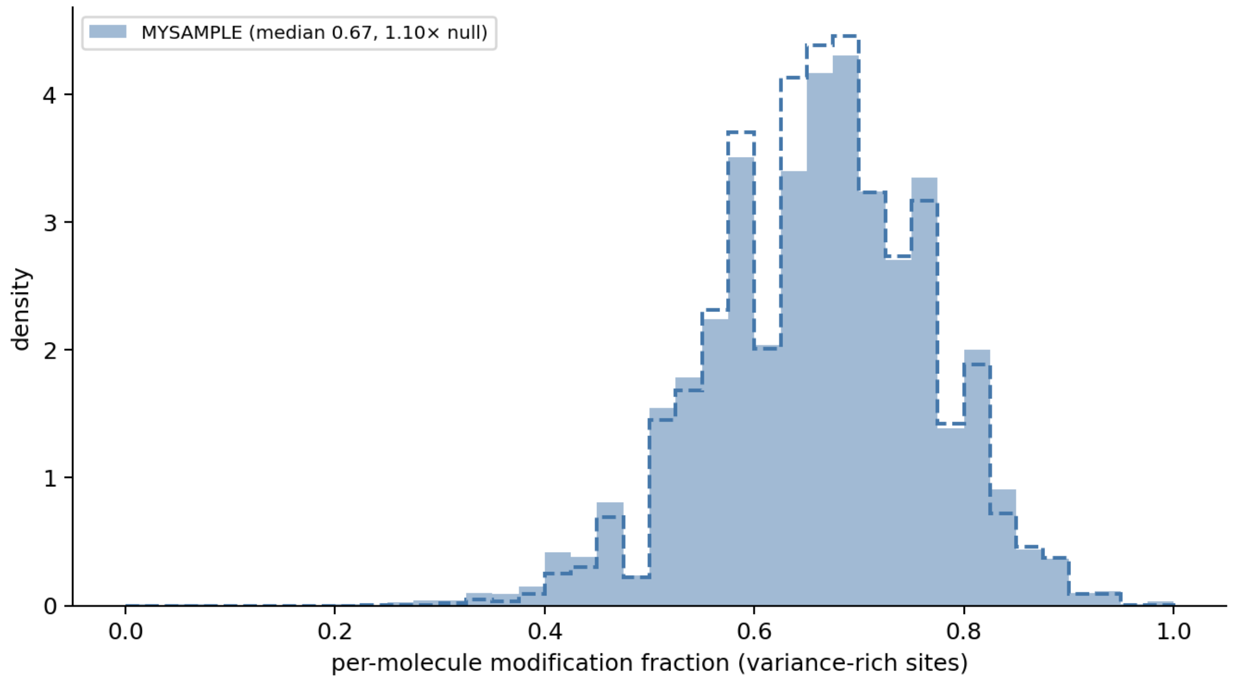


HeLa 28S rRNA

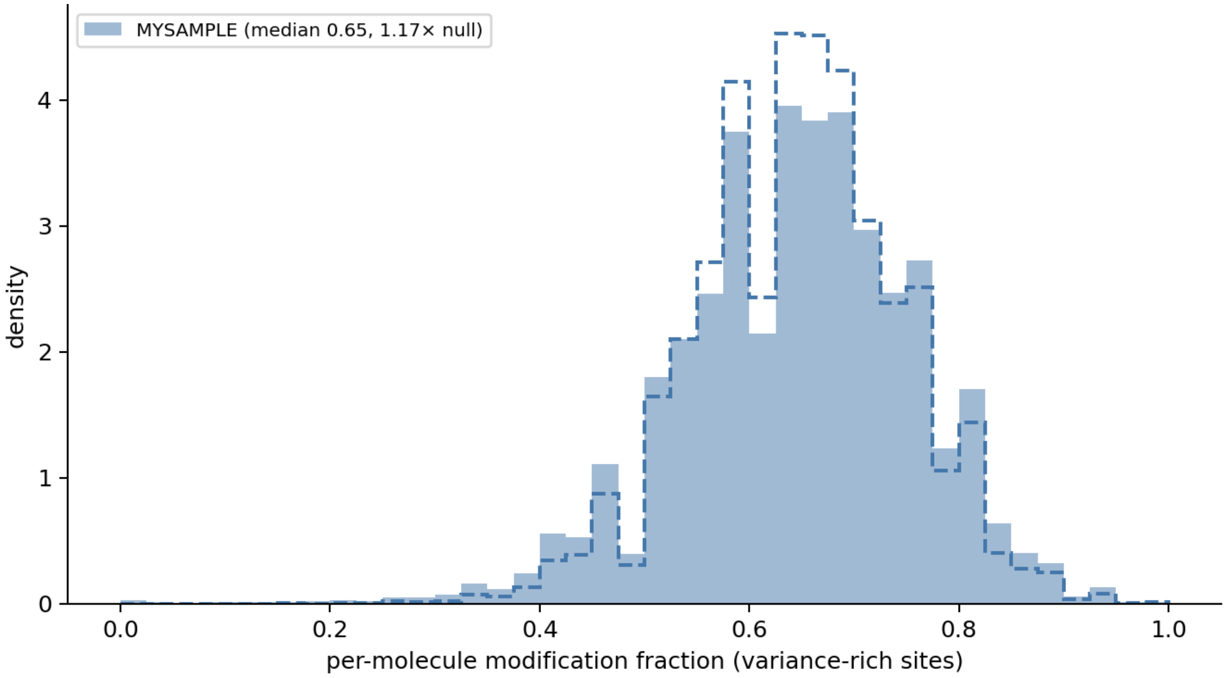


HCT116 18S rRNA

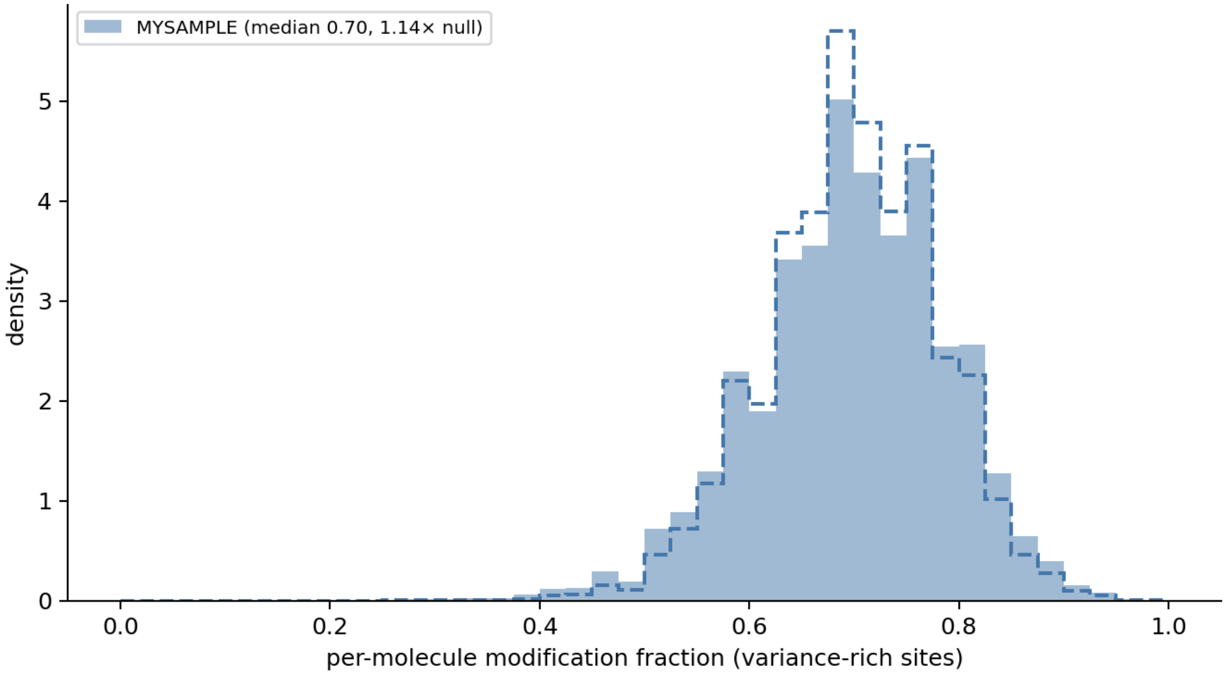


HCT116 28S rRNA

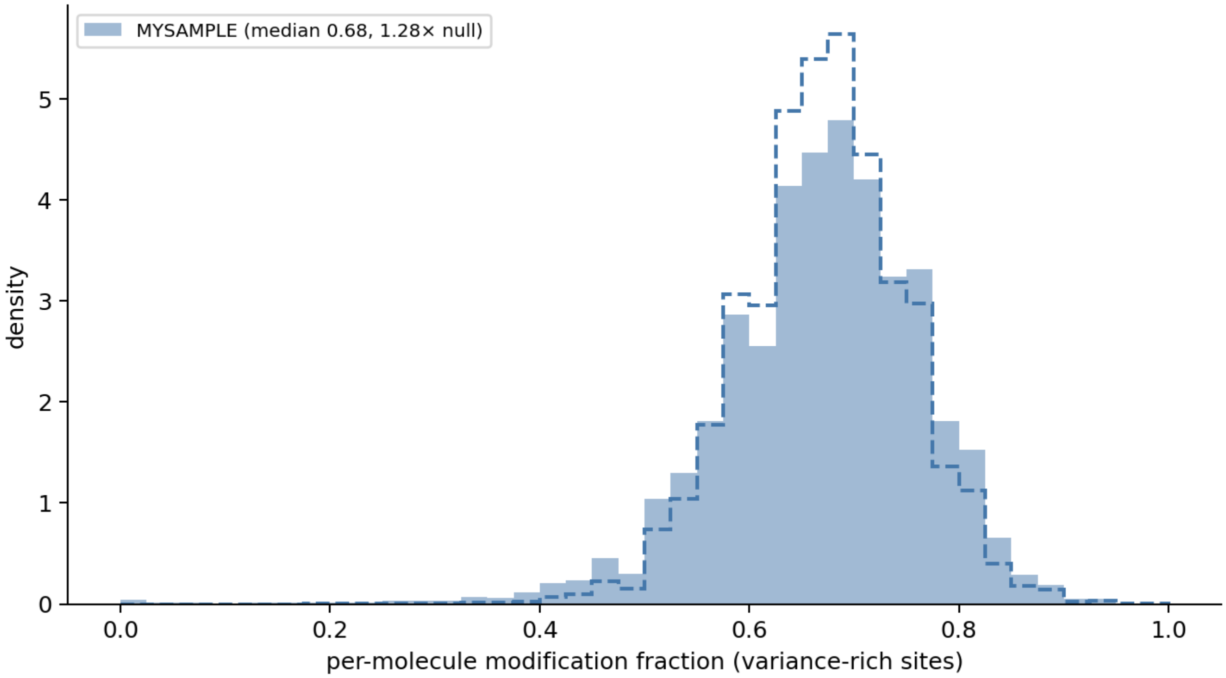
MYSAMPLE (median 0.68, 1.28× null)
density
per-molecule modification fraction (variance-rich sites)
0
1
2
3
4
5
0.0
0.2
0.4
0.6
0.8
1.0

**Figure S7.** Complete analysis of the 5.8S rRNA co-occurrence data set.

The 5.8S rRNA possesses 4 modifications, Um14, Ψ55, Ψ69, and Gm75. As stated in the text, Um14 is too close to the 5` end and is released by the helicase motor protein leading to fast translocation of this site through the nanopore sensor resulting in poor signal. The three remaining sites, which include Ψ55, Ψ69, and Gm75, illustrate the two artifacts that the method must control. All are modified at appreciable but very different fractional levels (0.46, 0.99, and 0.79, respectively, in HEK293T), and they lie 14, 6, and 20 nt apart. The signal for the Ψ69, at 0.99, is effectively saturated, in which only 143 of 15,660 molecules are unmodified at this position. Its apparent co-occurrence with Gm75 is high striking (OR = 3.13, 95% CI 2.29–4.28); however, these rest entirely on those 143 reads (panel A). This example illustrates how near-saturation inflates the odds ratio while the absolute deviation from independence remains negligible. Thus, Ψ69 falls above the near-saturation ceiling and is excluded from analysis by the criteria established above. Additionally, the Ψ55–Ψ69 and Ψ69–Gm75 pairs lie within the 20-nt neighbor-interference zone and would be excluded from the analysis for this reason, too.

Only one 5.8S pair, Ψ55–Gm75 at 20 nt, satisfies both criteria of having non-saturating occupancy and outside the interference zone, and the results show no evidence of co-occurrence (OR = 1.05, 95% CI 0.975–1.130; panel B). The 5.8S rRNA provides no interpretable co-occurrence signal because its three analyzable sites are too few, too clustered, and in one case too saturated.

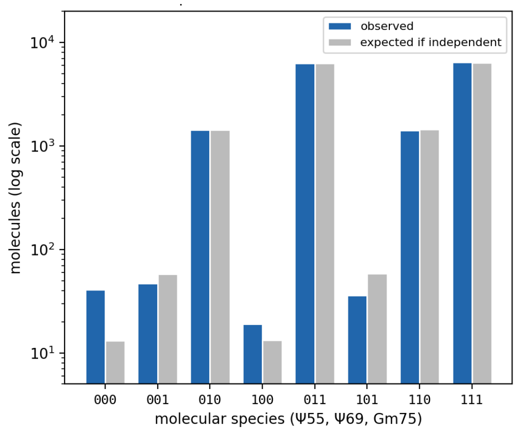


Panel A. Landscape of the three 5.8S rRNA modification pairs.

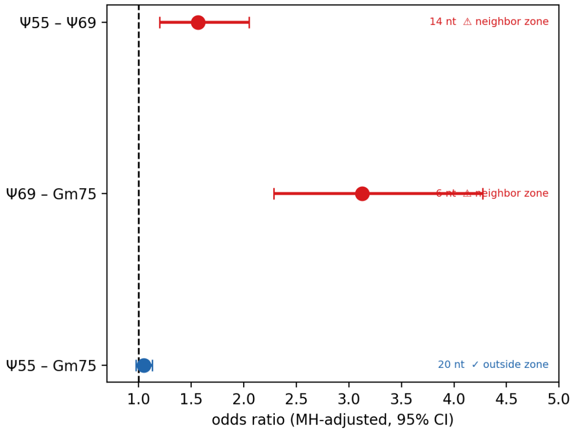
